# The evolution of competitive ability for essential resources

**DOI:** 10.1101/804542

**Authors:** Joey R. Bernhardt, Pavel Kratina, Aaron Pereira, Manu Tamminen, Mridul K. Thomas, Anita Narwani

**Affiliations:** Aquatic Ecology Department, Eawag, Überlandstrasse 133, CH-8600 Dübendorf, Switzerland; School of Biological and Chemical Sciences, Queen Mary University of London, Mile End Road, London E1 4NS, United Kingdom; Department of Biology, University of Turku, Natura, University Hill, 20014 Turku, Finland; Centre for Ocean Life, DTU Aqua, Technical University of Denmark, Kongens Lyngby, Denmark

**Keywords:** Chlamydomonas reinhardtii, coexistence theory, competition, eco-evolutionary dynamics, phytoplankton, R^*^, resource competition theory

## Abstract

Competition for limiting resources is among the most fundamental ecological interactions and has long been considered a key driver of species coexistence and biodiversity. Species’ minimum resource requirements, their R^*^s, are key traits that link individual physiological demands to the outcome of competition. However, a major question remains unanswered - to what extent are species’ competitive traits able to evolve in response to resource limitation? To address this knowledge gap, we performed an evolution experiment in which we exposed *Chlamydomonas reinhardtii* for approximately 285 generations to seven environments in chemostats which differed in resource supply ratios (including nitrogen, phosphorus and light limitation) and salt stress. We then grew the ancestors and descendants in common garden and quantified their competitive abilities for essential resources. We investigated constraints on trait evolution by testing whether changes in resource requirements for different resources were correlated. Competitive abilities for phosphorus improved in all populations, while competitive abilities for nitrogen and light increased in some populations and decreased in others. In contrast to the common assumption that there are trade-offs between competitive abilities for different resources, we found that improvements in competitive ability for a resource came at no detectable cost. Instead, improvements in competitive ability for multiple resources were either positively correlated or not significantly correlated. Using resource competition theory, we then demonstrated that rapid adaptation in competitive traits altered the predicted outcomes of competition. These results highlight the need to incorporate contemporary evolutionary change into predictions of competitive community dynamics over environmental gradients.

## Introduction

Resource limitation and competition for limiting resources are among the most important drivers of population growth [1], species distributions [2,3] and biodiversity [4]. Resource competition theory (RCT, [1]) predicts that a few key resource traits, including the minimum resource level a population requires to maintain positive population growth (R^*^), determine the outcome of competition over short time scales [5]. However, we still do not know how these resource traits evolve as populations adapt to new environments, especially in the context of organisms competing for essential resources such as light and nitrogen. This is an important gap in knowledge because rapid evolution may be able to alter competitive outcomes among species [6,7]. Understanding how evolutionary processes influence species’ traits that are relevant to coexistence is therefore critical to understanding the ecological mechanisms that create and maintain biodiversity [8]. Evolutionary change in one or multiple competing species can increase the likelihood of coexistence by reducing differences in species’ competitive abilities for a given resource (i.e. reducing ‘fitness differences’) and by altering the identity of the resource that each species finds most limiting (i.e. increasing ‘niche differences’) [9]. Since we do not currently understand the potential constraints on the adaptation of essential resource-use traits, we cannot predict the degree to which evolution contributes to or prevents competitive coexistence.

Resource competition often acts as a strong selective agent that drives patterns of biodiversity and trait change via character divergence [10,11] and adaptive radiation [12]. Competition can select for individuals that are able to consume ‘alternative’ resources, or those that are not shared with other competitors [13]. Over time, this results in adaptive trait divergence and niche differentiation [9,14]. Less well appreciated is that when resources are essential, or non-substitutable, opportunities for niche differentiation are limited and competition cannot be avoided by character displacement because all competitors require the same limiting resources [7,15]. In this case, selection favours improved competitive ability, or a reduced population-level R^*^ for the shared limiting resource [15,16]. However, adaptation may be constrained by physiological limits, genetic correlations between multiple traits [17], or lack of genetic variation in resource traits [18]. These constraints may be particularly strong in the case of adaptation to essential resource limitation because there are few opportunities for divergence in adaptive strategies.

Trade-offs among species in competitive abilities for different resources have been observed at large evolutionary scales (i.e. across clades) [19,20]. Turnover in species abundances across gradients of resource ratios suggests that these trade-offs underlie species distributions and patterns of biodiversity [1,21]. These trade-offs may arise due to differences in the local conditions in which the traits evolved, or from biophysical or genetic constraints that prevent individuals from optimizing several resource-use traits simultaneously. There are at least two types of trade-offs which can govern resource competition: gleaner-opportunist trade-offs (Figure 1A), (Electronic Supplementary Material (ESM) Figure S1) [22,23], and trade-offs in the ability to acquire different essential limiting resources (e.g. light versus nitrogen or nitrogen versus phosphorus; Figure 1B) [19,20,24,25]. A gleaner-opportunist trade-off is a trade-off between a low minimum resource requirement and a high maximum growth rate. A gleaner phenotype grows better at low resource levels and an opportunist phenotype can take advantage of high resource levels [26] (Figure 1A). Although the existence of trade-offs in resource-use traits has been demonstrated on a macroevolutionary scale spanning large swaths of evolutionary time, the microevolutionary processes by which they may arise and the mechanisms that maintain them are still poorly understood.

**Figure 1.**
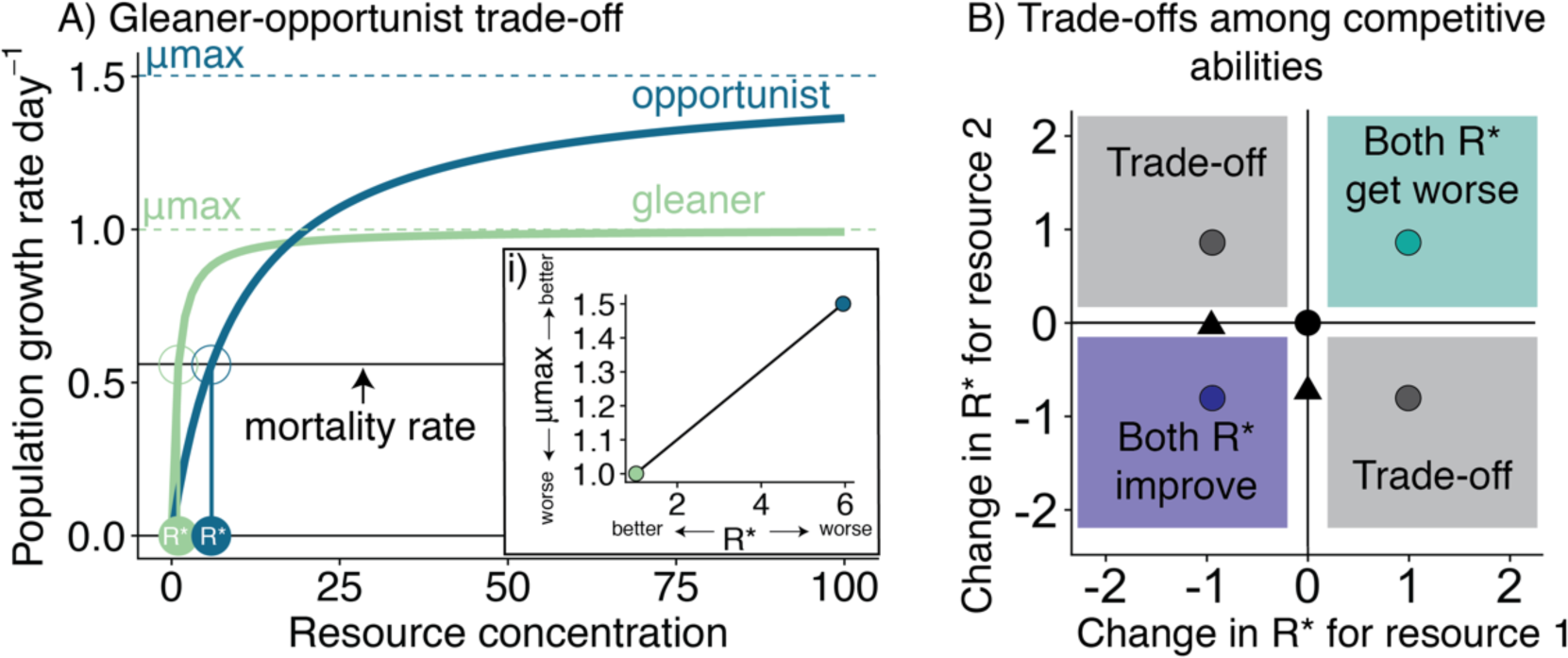
A) Two example Monod curves [27], describing resource-dependent population growth rates demonstrate a gleaner-opportunist trade-off (i.e. a trade-off between individuals that have high growth rates at low resource levels (green curve, gleaner), and lower growth rates at high resource levels compared to opportunists (blue curve, opportunist)). R*, shown in circles, are the resource concentration at which population growth rate is zero. Here we show a mortality rate of 0.56/day, consistent with the dilution rate in our experiments. A gleaner-opportunist trade-off may be detected empirically by a positive relationship between μmax and R* (panel A, inset i). B) Trade-offs may arise when adaptation to one environment comes at the cost of performance in a different environment (e.g. grey dots in grey regions), here shown in terms of changes in R* of descendant populations relative to their ancestors (black dot in centre). Alternatively, adaptation may arise via improvement in multiple traits simultaneously (e.g. purple dot, lower left quadrant), or conditional neutrality (i.e. improvements in one trait dimension, but no cost in another, black triangles). Maladaption may occur if there are losses of performance in multiple traits simultaneously (turquoise dot, upper right quadrant).

Ecological and evolutionary trade-offs are expected to arise from fundamental constraints on the use and acquisition of energy and materials. Organisms have fixed resource-and energy budgets with which to metabolize, grow and reproduce, such that energy and resources allocated to performing one function necessarily cannot be used for performing another independent function [28,29]. Furthermore, the observation that no single genotype or phenotype maximally performs all functions necessarily implies that there are physiological constraints preventing the evolution of “Darwinian demons” [30]. Despite the fact that evolving individuals eventually will face trade-offs, not all local adaptation comes at a cost. First, trade-offs may not occur when multiple functions can be optimized using the same energetic and resource allocations. For example, this may occur when metabolic pathways affecting multiple functions are highly connected and interdependent. Increasing efficiency in any part of the metabolic pathway may therefore also reduce demands in the rest of the network [31]. In phytoplankton, this may be the case for resource requirements for light and nitrogen because chloroplasts are typically very nitrogen-rich [32,33]. Similarly, proteins required for nutrient uptake and metabolism are produced by phosphorus-rich ribosomes [32,33]. Second, trade-offs between competitive abilities for different resources may not arise if local adaptation results in energy and material budgets that are larger overall - i.e. they are still approaching a fitness optimum. Finally, mutations that improve fitness in a local environment may result in trade-offs due to antagonistic pleiotropy [28] or in mutation accumulation for traits that are not under selection [34]. However, other outcomes are also possible, including neutral genetic variation or synergistic pleiotropy [35,36]. Evidence that pleiotropy and mutation accumulation should consistently generate trade-offs (rather than fitness-neutral or positive trait change in an alternative environment) is still lacking [28].

To understand how essential resource competition traits evolve and how adaptation is constrained, we used experimental evolution with a model organism, *Chlamydomonas reinhardtii*. Experimental evolution allowed us to control the ecological conditions of selection in chemostat, to isolate the effect of single limiting resources, and to minimize confounding selective forces across treatments and replicates. We created seven distinct selection environments in chemostats that varied either in the supply of essential resources or salt concentration and quantified how populations’ resource-competition traits and salt tolerances evolved. We replicated the evolutionary treatments across five ancestral populations in order to quantify heterogeneity in the responses to selection, and the repeatability of evolutionary outcomes [37]. Using whole genome resequencing of the ancestors and descendants of the evolution experiment, we confirmed that the descendants had fixed mutations over the course of the experiment, and were no longer genetically identical to the ancestors, suggesting that the observed phenotypic changes have a genetic basis.

We tested three predictions:

1. When populations are exposed to limitation of essential resources, selection on resource-use traits should reduce R^*^, the minimum resource requirement. Additionally, evolutionary changes in R^*^ should be larger in the genotypically diverse population relative to the isoclonal populations [38] because adaptation from standing genetic variation can occur more rapidly [39] than adaptation acting on novel mutations [40]. Lastly, we predicted that salt stress, in addition to resource limitation would lead to greater adaptive trait change, particularly because stress can increase rates of mutation [41].
2. Adaptive trait change is subject to trade-offs. Trade-offs between competitive abilities for different resources, gleaner-opportunist trade-offs, or trade-offs between resisting salt stress and having a high growth rate or low R^*^ may constrain or structure adaptive change in resource traits [16,20] and potentially cause adaptation in one environment to come at a cost to performance in another environment [42] (Figure 1B). Alternatively, positively correlated competitive traits may cause selection for a lower R^*^ for one resource to reduce R^*^ for another (pleiotropic or correlated fitness benefits in low-resource environments) [42–44].
3. We predicted that if trade-offs in resource-use traits cause traits to diverge across different selection environments, this would increase the chance that populations selected in different environments could competitively co-exist.

## Methods

### Evolution experiment

We obtained a strain of *C. reinhardtii* (cc1690 wild type mt+) from the Chlamydomonas Center (chlamycollection.org). We selected four colonies derived from single cells (hereafter referred to as Anc 2, Anc 3, Anc 4 and Anc 5, ESM Appendix B Figure S16), and inoculated them into liquid COMBO freshwater medium [45]. We randomly assigned seven small chemostats (28 mL) to each of the four isoclonal ancestral populations (Anc 2-5) and the genotypically diverse population, cc1690. The seven chemostats assigned to each of the ancestral populations were then randomly assigned to one of seven treatments which we maintained for 285 days: COMBO (hereafter referred to as C), nitrogen limitation (N), phosphorus limitation (P), light limitation (L), salt stress (S), biotically-depleted medium (i.e. medium previously used to grow seven other species of phytoplankton, which was then filtered and sterilized) (B), and a combination of salt stress and biotically-depleted medium (‘BS’). The C treatment had COMBO medium supplied with an equable resource ratio (i.e. not highly limited in a single nutrient), which allowed us to compare specific adaptations to resource-limitation to adaptations to life in chemostat more generally. Here we used the term ‘population’ to refer to Anc 2, Anc 3, Anc 4, Anc 5, cc1690 (the ‘ancestors’) as well as all of their descendant populations (‘descendants’). In total, there were five ancestral populations, and 32 descendant populations because three were lost to contamination. Detailed information on experimental evolution methods is available in the Supplementary Methods in the ESM (Appendix A).

### Determination of R^*^ and salt tolerance

We determined the minimum resource requirements for positive population growth (R^*^) for each population [1] via batch culture experiments. We defined N^*^ as the minimum nitrogen concentration and P^*^ as the minimum phosphorus concentration for positive population growth. We define I^*^ as the minimum light level required for positive population growth (similar to I_c_ in [46]). We estimated R* by measuring population growth rates at ten resource levels for each of nitrogen, phosphorus and light for three days (see ESM Appendix A: Supplementary Methods: *‘Competitive trait assays’* for more details on the resource levels, acclimation and measurements). We estimated ‘consumption vectors’ ([1]) for N and P via stoichiometry of exponentially growing populations [3], and cell size by measuring single cell lengths using a high throughput imager (Biotek® Cytation 5), and calculating cell biovolume assuming cells were spheres using 4/3 × π × radius^3^.

In order to determine populations’ R*, we modeled resource-dependent population growth via a Monod curve [1,35]. We estimated the parameters of the Monod curve directly from population-level time series of chlorophyll-*a* relative fluorescence units measured over the resource gradients. We modeled the resource-dependent rate of population growth, *r*, during the exponential phase as:

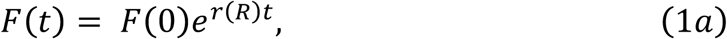

where *F*(*t*) is the population-level RFU at time *t*, and *r*(*R*) is given by:

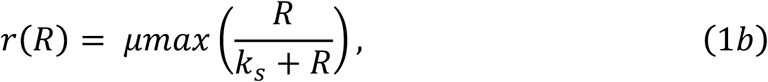

using nonlinear least-squares regression with the *nls.LM* function in the *minpack.LM* package [36] in R. Population growth rate, *r*, is a function of *µmax*, the maximum population growth rate, *R*, the resource concentration, and *k*_*s*_, the half-saturation constant for population growth.

Using the estimated parameters of the Monod curve (i.e. Equation 1b), we estimated R^*^ as:

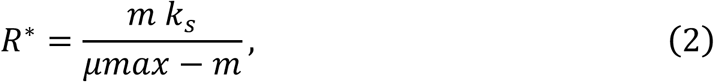

where *m* is the mortality rate, which we set to be 0.56/day to reflect the mortality caused by dilution in chemostat experiments. To simplify our analyses, we used Equations 1 and 2 to estimate minimum light requirements (I^*^), where *R* = irradiance. We also included ESM Figure S2 with parameters estimated from an Eilers-Peeters curve [37] for light.

To estimate the uncertainty in the Monod curve (Equation 1) fits, we determined confidence intervals around the fitted Monod curves using non-parametric bootstrapping of mean-centered residuals using the *nlsBoot* function with 999 iterations in the *nlstools* [38] package in R. We calculated 95% confidence intervals as the range between the 2.5th and 97.5th quantiles.

We defined the salt tolerance as salt concentration at which growth rates are half their maximum (which occurs at a salt concentration of zero). We estimated salt tolerance by modeling population growth rates during the exponential phase, *r*, as a function of salt concentration, *S*, using a simplified form of the logistic function:

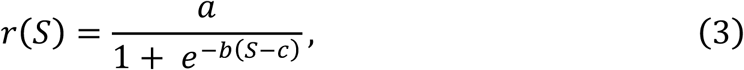

where *a* (the upper asymptote), is the maximum population growth rate (not salt-stressed), *b* is the decline in growth rate with increasing salt concentration and *c* is the salt concentration at which growth rates are half their maximum, in g/L.

### Quantifying trait change and testing for trade-offs

We tested for changes in R^*^ between descendant and ancestral populations by subtracting the ancestral trait value from the descendant trait value and quantifying whether the 95% on the difference overlapped zero. We tested whether the change in resource-use traits was greater in the genotypically diverse populations than the isoclonal populations by comparing the 95% CI of the trait changes.

We tested for trade-offs between:

1. growth rates at high vs low supply of a given resource (ie. µmax vs R^*^, or a gleaner-opportunist trade-off) (Figure 1A),
2. competitive abilities for different resources, or competitive ability and cell size, and
3. changes in traits in multiple dimensions (Figure 1B).

We tested for trade-offs using multiple linear regressions. We quantified competitive ability for a given resource as 1/R^*^ [20], and tested for trade-offs among competitive abilities for different resources (trade-off 2). In order to assess trade-offs among multiple traits and cell size, we centered and scaled the variables using the *scale* function in R (mean=0, standard deviation=1) so all variables could be compared on the same scale. In all cases of multiple regression, we included ancestor ID as a fixed effect to account for relationships among ancestors and descendants.

We tested for differences in multivariate trait change as a function of selection treatment and ancestor using redundancy analysis (RDA) with the *capscale* function in the R package *vegan* [50], version 2.5-4. Here we included all of the traits we measured: R^*^s, cell biovolume, consumption vectors (i.e. stoichiometry), and salt tolerances. We used permutation tests (*anova.cca* in *vegan*) to test the statistical null hypothesis that selection treatment and ancestor ID had no significant impact on any independently varying linear combination of traits. We used the same approach to test the effects of treatment on trait variation along the individual axes. We assessed which descendant populations had diverged from their ancestors in different environments using post-hoc Tukey tests using the *TukeyHSD* function in R. We conducted all of our statistical analyses using R, version 3.6.1 [51].

### Quantifying genetic changes associated with selection environments

DNA was extracted using a chloroform-methanol extraction and libraries were prepared using the Bioo Scientific NEXTflex Rapid Illumina DNA-Seq Library Prep Kit. For details and bioinformatic methods, please refer to the Supplementary Methods in the ESM (Appendix A).

### Testing the potential for altered predicted outcomes of competition

We used resource competition theory [1] to predict the outcome of pairwise competition for two resources: nitrogen and phosphorus. RCT predicts that two populations can coexist stably if they meet three conditions: 1) their zero net growth isoclines (ZNGIs) cross (i.e. populations differ in the identity of the resource that most limits their growth), 2), they each consume more of the resource which most limits their growth (i.e. each population has a steeper consumption vector for the resource which is most limiting to it) and 3), the supply point of resources in the environment falls above their ZNGIs and between the consumption vectors of the two populations. If the pair of populations meets criterion 1 and 3 but not 2, theory predicts unstable coexistence or priority effects. If the pair of populations meets one or none of these criteria, theory predicts competitive exclusion. We compared all possible combinations of the five populations in their ancestral state and after selection in the different resource environments. We then assessed the proportion of these pairwise interactions that would be expected to lead to unstable coexistence, stable coexistence or competitive exclusion.

## Results

### Evolutionary changes in R^*^ and salt tolerance

Relative to their ancestors, P^*^ declined in all five populations exposed to P limitation (P, Figure 2A). Declines in P^*^ ranged from 43% to 85% across the replicate populations. In response to N limitation, N^*^ declined in two populations (14%, 34% decline), did not change in two populations and increased in one population (47% increase) (N, Figure 2B). I^*^ increased in two populations exposed to low light (L, 12%; 28% increase), and did not change in the remaining three populations exposed to low light (Figure 2C). Salt tolerance increased in all populations exposed to high salt (93% - 369% S and BS, Figure 2D). Consumption vectors, quantified as the P:N molar ratio in the biomass of populations growing exponentially, decreased in all of the populations subjected to nitrogen limitation, and increased in four of the five populations exposed to phosphorus limitation (ESM Figure S4, see also Figures S5-6). This suggests that populations selected under nitrogen limitation contained more nitrogen relative to phosphorus whereas populations selected under low phosphorus contained more phosphorus relative to nitrogen. Contrary to our predictions, the descendants of the genotypically diverse cc1690 population did not show more trait change than any of the isoclonal populations (triangles vs small dots in Figure 2). However, the genotypically diverse cc1690 population did match our predictions in terms of the direction of adaptive trait change in all selection environments: P^*^ decreased under P-limitation, N^*^ decreased under N-limitation, salt tolerance increased in the high salt environment, and I^*^ decreased under low light, though the change in I^*^ was not statistically significant.

**Figure 2.**
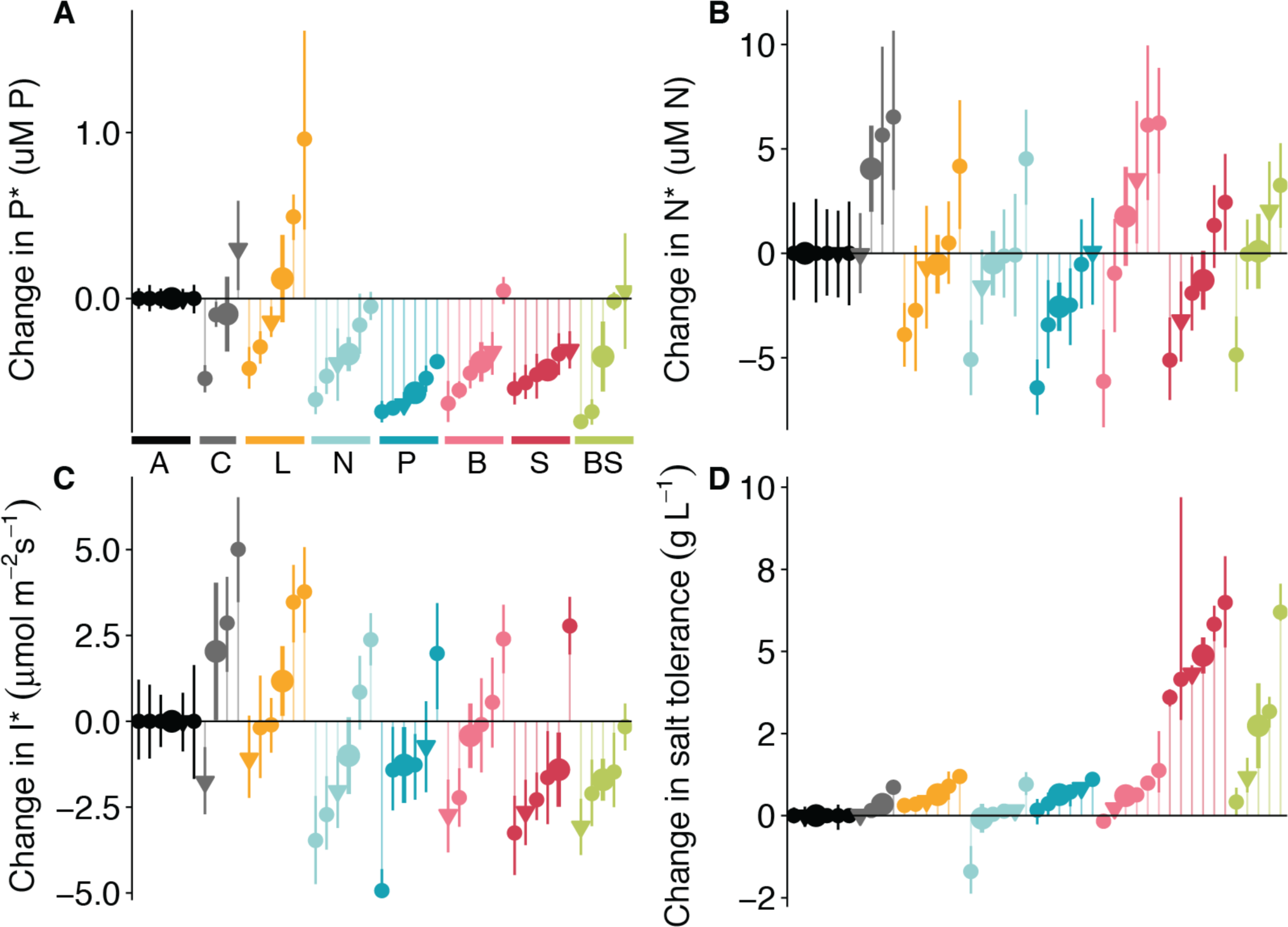
Changes in resource competition traits relative to ancestors in seven different selection environments (A: ancestors, C: COMBO, L: light-limited, P: P-limited, N: N-limited, B: biotically depleted media, S: high salt, BS: biotically depleted and high salt). Small points correspond to individual populations and large points correspond to the average change (error bars are ± 1 SE) of all populations in a given environment. Populations represented with a triangle are the genotypically diverse populations (cc1690), circles are isoclonal. Error bars on small points in A-D correspond to 95% CI from non-parametric bootstrapping. Colour legend in A applies to all panels.

When considering all traits together, descendant populations diverged from their ancestors, and variation in these new phenotypes was associated with selection environment (Redundancy Analysis, Figure 3). We tested for constraints on adaptive change by assessing whether there was significant separation between ancestors and descendants on the RDA axes. RDA axes 1 and 2, which represent linear combinations of selection environment and ancestor ID (PERMANOVA p < 0.01) explain 85% of the variation associated with selection environment, and 36% of the total variation. On RDA axis 1 (PERMANOVA p < 0.001), which is primarily associated with variation in salt tolerance and P*, populations selected in the P, S, B and BS environments were significantly different (separated) from the ancestors. The salt selected populations (S and BS) were also different from the COMBO (C) and low-light selected treatments (L). On RDA axis 2 (PERMANOVA p = 0.005) which is associated with variation in P:N (consumption vector slope), P is different from ancestors and the C, L, B, S, BS populations. The RDA showed that most of the variation in multivariate phenotypes across selection environments was associated with variation in salt tolerance and P*, and much less independent variation was associated with N* and I* (Figure 3), suggesting that variation in these traits may be subject to physiological or genetic constraints.

**Figure 3.**
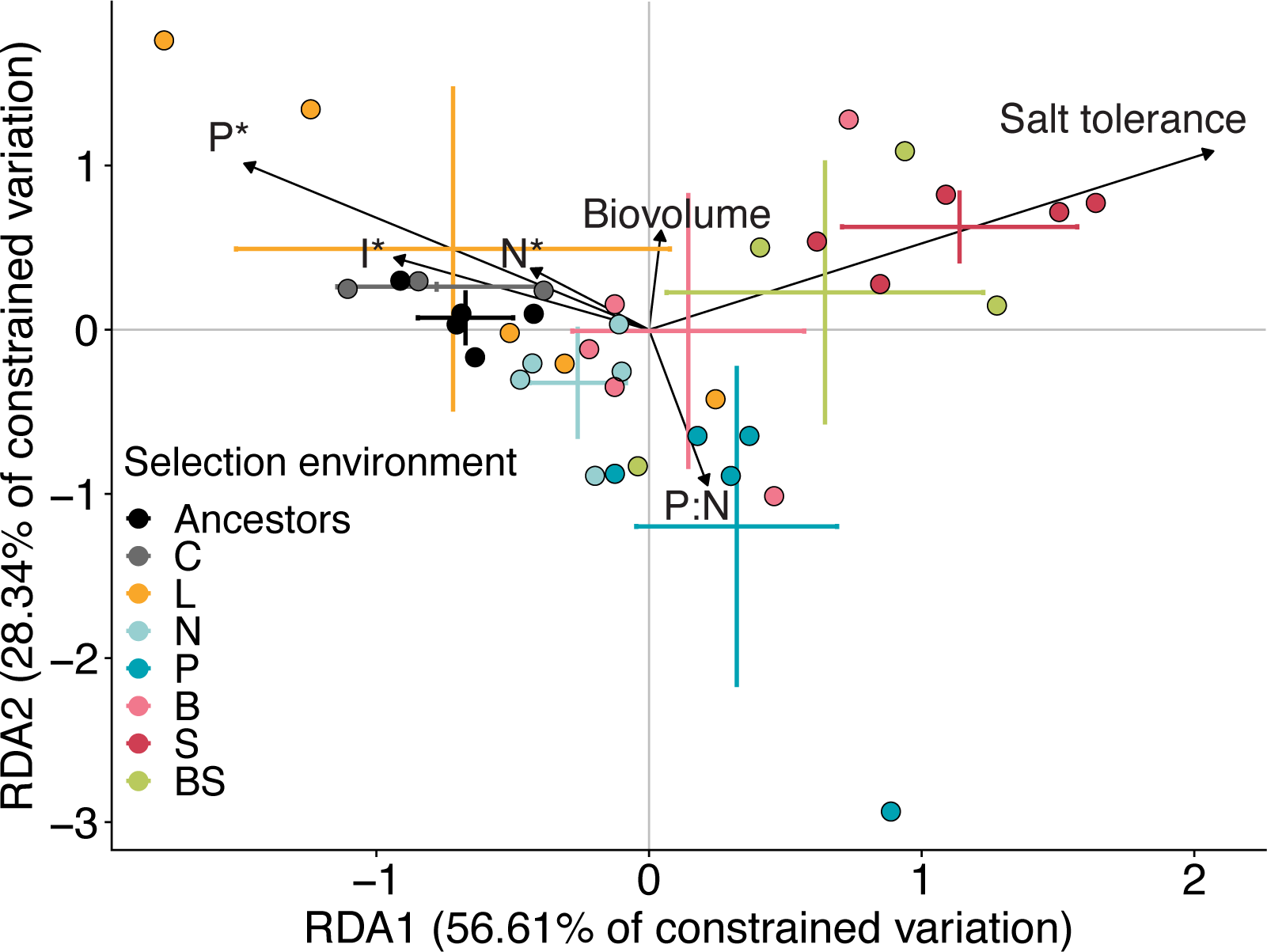
Redundancy analysis of N*, I*, P*, salt tolerance, consumption vectors and cell biovolume across selection environments. Error bars correspond to standard error around treatment means (n = 5 per treatment). RDA axes 1 and 2 (PERMANOVA p < 0.01) explain 85% of the variation associated with selection environment, and 36% of the total variation.

### The structure of trait variance: observed trade-offs

Maximum growth rate across populations increased with minimum resource requirements (R^*^) for light (OLS slope = 0.021, 95% CI: 0.0027, 0.039, Adjusted *R*^*2*^ = 0.11), for nitrogen (OLS slope = 0.019, 95% CI: 0.0078, 0.031, Adjusted *R*^*2*^ = 0.30), and for phosphorus (OLS slope = 0.12, 95% CI: 0.0094, 0.23, Adjusted *R*^*2*^ = 0.13), indicating a trade-off between growth at high and low resource supplies (because a lower R^*^ indicates faster growth at minimum resource levels) (Figure 4, A-C). Across populations, competitive abilities for N and P (CN and CP) were positively associated (ESM Tables 4-6, Figure 4D, ESM Figures S13-15). After accounting for covariance with competitive abilities for other resources and ancestor ID, competitive abilities for light were negatively associated with cell biovolume, while and N and P competitive abilities were not related to cell size (ESM Tables 4-6, ESM Figures S13-15). Principal components analysis of cell biovolume, competitive abilities for light, nitrogen and phosphorus showed that 74% of the variation in cell volume and competitive abilities is explained by the first two PC axes. The first two PC axes demonstrate a positive association between competitive abilities for N and P, and a possible trade-off between biovolume and competitive ability for light (Figure 4D).

**Figure 4.**
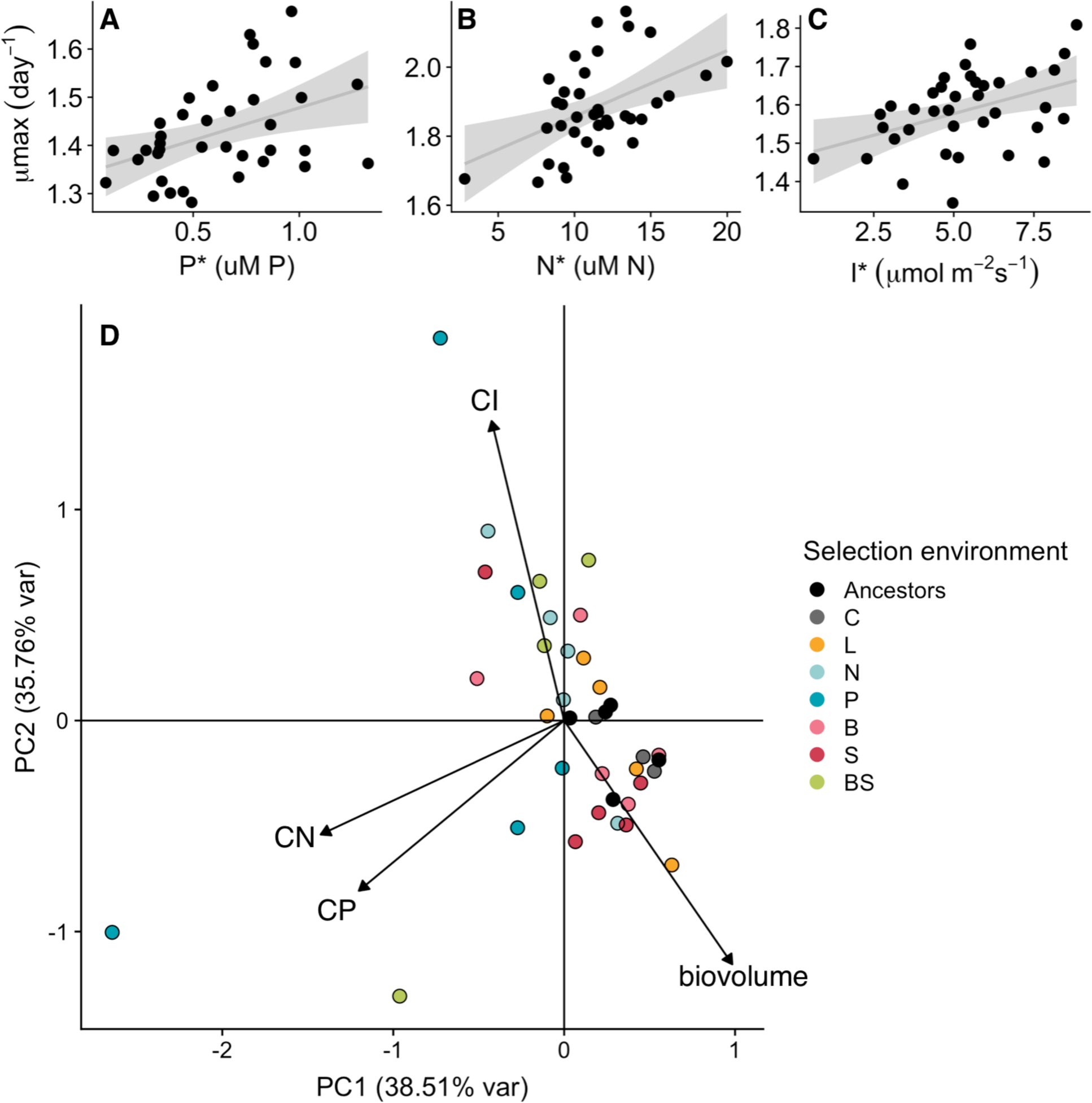
High maximum population growth rates (µmax) are positively associated with high minimum resource requirements for phosphorus (A), nitrogen (B) and light (C). Estimates of µmax and R^*^ in A-C are from Monod curves generated over independent gradients of phosphorus, nitrogen and light. D) Loadings on the first two PC axes of a PCA using competitive abilities for phosphorus, nitrogen and light (CP = 1/P^*^, CN = 1/N^*^, CI = 1/I^*^) and cell biovolume. 74.3% of the variation in resource use traits and cell size is explained by the first two PC axes.

### Correlations in changes across traits

Though theory often assumes that competitive abilities for different resources are negatively related [16,52], our results did not support this finding either when considering absolute variation in competitive abilities (ESM Tables 4-6) or variation in the change in R^*^ relative to the ancestral populations (Figure 5; ESM Tables 1-3, ESM Figure S10). The changes in R^*^ for different resources never showed evidence of any trade-offs, and instead were either positively associated (Figure 5A, B) or showed no significant relationship (Figure 5C, ESM Figure S10).

**Figure 5.**
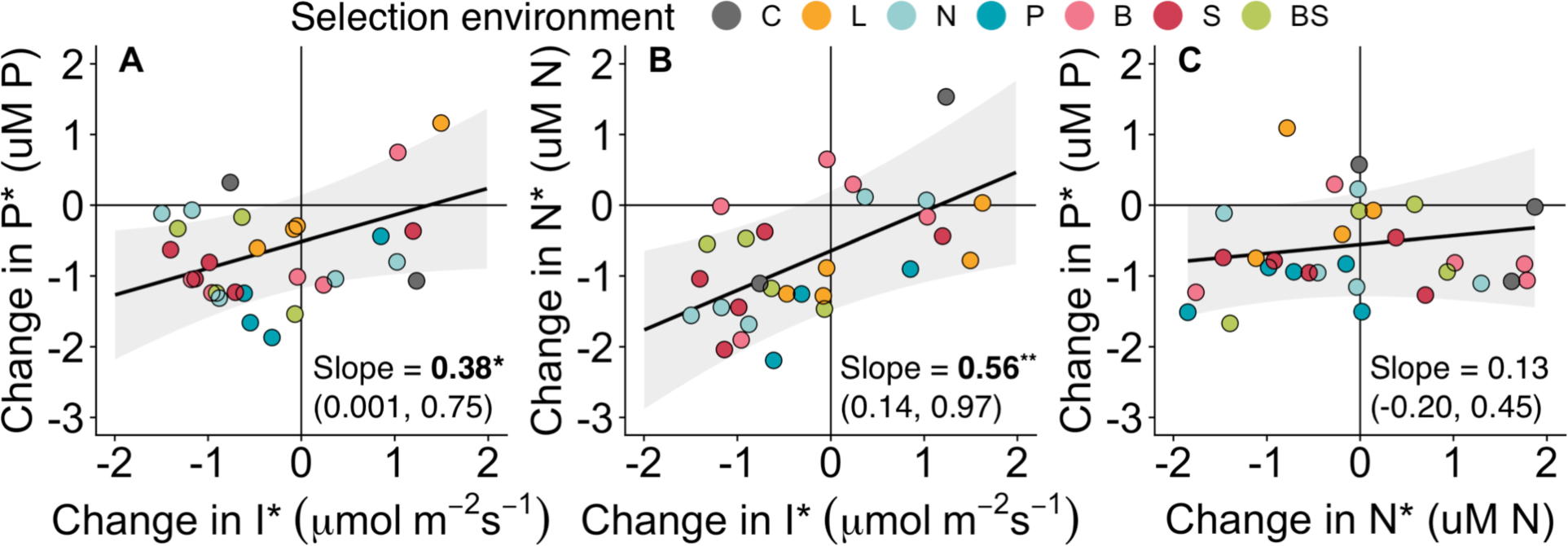
Partial regression plots showing how changes in descendants relative to ancestors in two traits are related to each other, while holding all factors in the statistical model that are not being displayed constant (complete model results in ESM Tables 1-3). Positive slopes indicate positively associated trait changes.

### Genetic changes after selection

Genetic differences between the ancestral and descendant populations were identified by whole genome re-sequencing. The presence or absence of single nucleotide polymorphisms (SNPs) identified within the populations were compared between the ancestors and descendants for each selection treatment. The number of variable SNPs ranged from 396 to 582 (ESM Table 7), roughly corresponding to mutation fixation rates of 1.25e-8 to 1.83e-8 mutations/[locus × generations]. Contrary to our expectations, salt stress did not increase the number of fixed mutations. Selection treatment had no significant effect on the total number of fixed mutations (ESM Figure S11 A; ANOVA p = 0.788), but the effect of the ancestor was highly significant (ESM Figure S11 B; ANOVA p < 1e-7).

### Evolutionary adaptation altered the predicted outcomes of competition

Divergence in minimum resource requirements and consumption vectors among populations of the same ancestor selected in different environments (Figure 3) was sufficient in some cases to lead to predicted coexistence. One such example is illustrated in Figure 6, where descendants of Ancestor 3 selected in low light and high salt environments have diverged sufficiently in their P^*^ and N^*^ such that they could possibly coexist. While neither descendant population could coexist with Ancestor 3, the two descendant populations could coexist with one another at a supply point illustrated as a yellow dot. In their ancestral state, out of all pairwise combinations of our five ancestor populations, resource competition theory [1] predicts unstable coexistence in four of 10 cases and competitive exclusion in five of 10 cases, and stable coexistence in one of 10 cases. After selection across the range of environments in our study, resource ratio theory predicts unstable coexistence in 19.9% of all possible pairwise interactions (698 total), stable coexistence in 27.94% of all possible pairwise interactions and competitive exclusion in 52.15% of all possible interactions (ESM Figure S12). Among populations selected in the same environment (59 total), resource ratio theory predicts competitive exclusion in 47.5%, stable coexistence in 32.2%, unstable coexistence in 20.33%.

**Figure 6.**
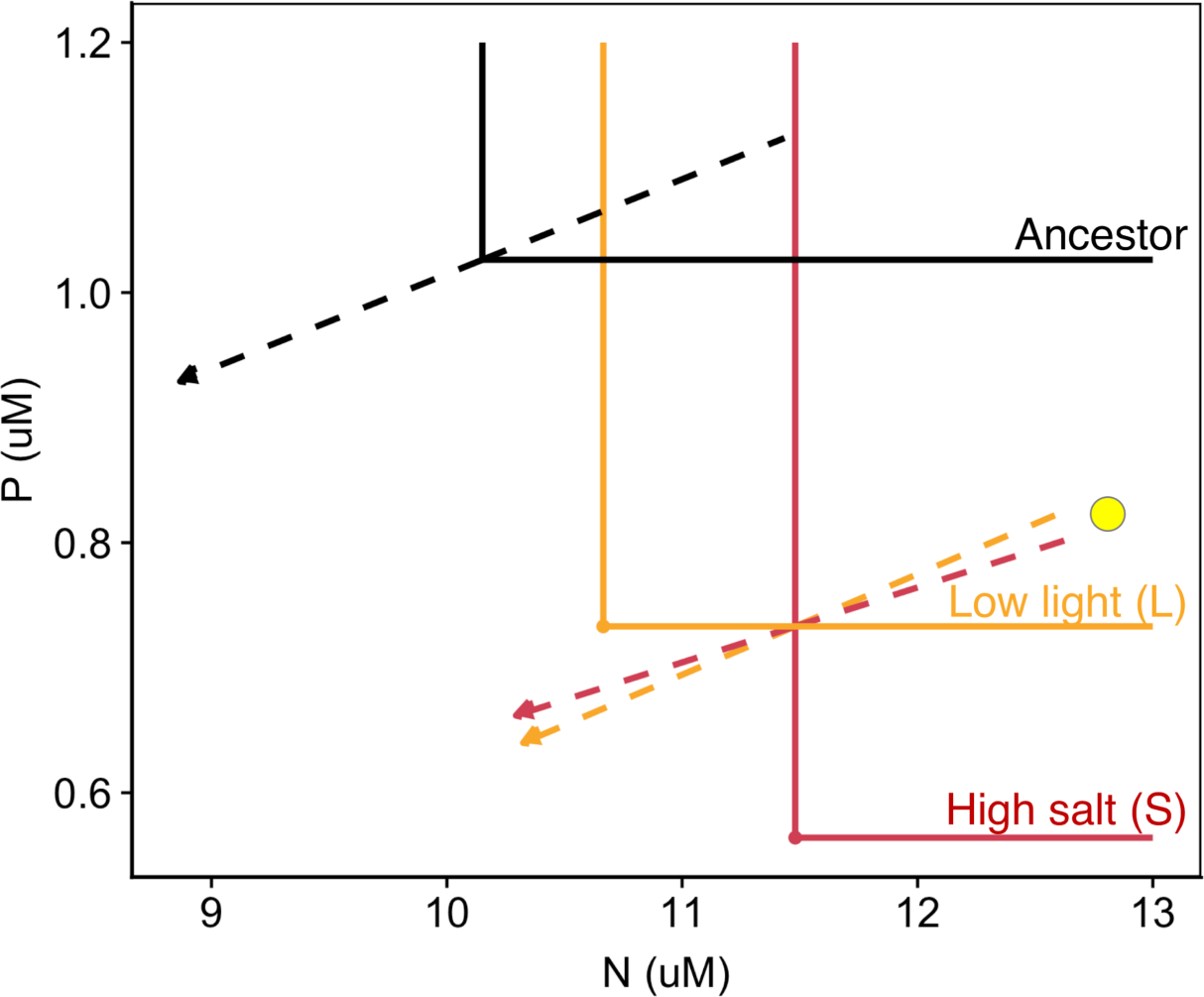
Descendants of Ancestor 3 evolved in high salt (red) and low light (orange) environments have diverged in their P^*^ and N^*^ such that they can co-exist. Neither descendant could co-exist with the ancestor. Zero net growth isoclines (solid lines) and consumption vectors (dashed lines) for Ancestor 3 (black) and the descendant of Ancestor 3 selected in low light (orange) and high salt (red).

## Discussion

Resource competition is among the most important processes structuring ecological communities [53], but competition theory often assumes that traits underlying competitive abilities remain fixed over ecological time scales [1,54]. Here we showed that the traits that underlie competitive abilities for essential resources can adapt rapidly in new resource-limited environments. Populations of *C. reinhardtii* often adapted to resource limitation by reducing their minimum resource requirements. When exposed to high salt, populations evolved higher salt tolerances. Not only could populations respond adaptively to new environments, but they could also adapt within approximately 285 generations. While we observed gleaner-opportunist trade-offs, we did not find evidence for trade-offs in competitive abilities for different resources.

Instead, adaptive changes in competitive ability for one resource were often positively associated with improvements in competitive ability for another. Since the ancestral and descendant populations were maintained under identical conditions when quantifying their traits, the changes we observed were heritable. We documented genetic changes as fixed single nucleotide polymorphisms in each of descendant lineages (ESM Appendix B Figure S16, Appendix C Table 7), which likely contributed to the heritable phenotypic changes we observed. Due to a lack of annotational information on the genes in which mutations were fixed, our ability to infer the connection between genotype and phenotype is limited. Future studies investigating the roles of gene expression regulation and epigenetic modification in contributing to heritable trait change of resource requirements would provide additional insights [55].

The magnitude of evolutionary change varied among resource competition traits. When considering all the populations together, adaptive change in P* was large (up to 85% decrease relative to ancestors), while adaptive change in N* was more limited, and change in I* was sometimes maladaptive. It is possible that the lack of adaptive change in N* and I* was because the ancestral populations in our experiment were already at or near a fitness optimum. Although there were no consistent differences in the magnitude of adaptive trait changes when comparing the genotypically diverse populations to the isoclonal populations, in all selection environments the direction of trait change was adaptive in the genotypically diverse populations. However, the absence of replicated genetically diverse populations within each treatment limits our ability to generalize the effects of genotypic diversity on evolutionary outcomes in a given environment.

Trade-offs in resource-use traits have been invoked to explain changes in dominance across supply ratio gradients and the coexistence of as many species as resources [1,19]. Trade-offs in competitive abilities for nitrogen and phosphorus [20], iron and light [24], and light and nitrogen [25] have been documented among and within species of phytoplankton. These trade-offs may arise due to local adaptation, or due to biophysical constraints on the acquisition or metabolism of different resources [19,52].

Individuals may invest resources into two main types of cellular machinery: uptake or assembly machinery. Uptake machinery is composed of nutrient uptake proteins and chloroplasts, which are both relatively nitrogen-rich, and both of which may scale with cell size because uptake and photosynthesis must take place at the cell surface. Assembly machinery, primarily composed of ribosomes, is relative phosphorus-rich and may also depend on cell size, as smaller cells tend to grow faster (“growth rate hypothesis” [33]). Consistent with expectations, competitive abilities for light were negatively associated with cell size, but in contrast to expectations, N and P competitive abilities were not. Furthermore, we did not find evidence for a trade-off between competitive abilities for nitrogen and phosphorus.

No evidence for trade-offs in competitive abilities for different resources is in contrast to observations of negative multivariate correlations observed on macroevolutionary timescales [20]. This runs counter to the idea that population genetic variation occurs along the same axes as variation among species - along ‘genetic lines of least resistance’ [56]. There are multiple potential reasons for this lack of observed trade-offs in competitive ability at this evolutionary scale. The first possibility is that essential resource requirements differ from other traits because they are linked via shared metabolic pathways in a metabolic network that controls the uptake, conversion and allocation of energy and materials. Requirements for different resources are intrinsically, metabolically linked and therefore non-independent. This suggests that observed trade-offs in R* at macroevolutionary scales are the result of major metabolic innovations across clades, breaking these metabolic linkages [57]. It is also possible that correlations observed at macroevolutionary scales are due to responses to local selection pressures that are unrelated to resource limitation, including grazing, disease and turbulent mixing [58]. A third possible explanation is that the descendant populations in our experiment had not yet reached fitness or trait optima, and as such, continued adaptation did not impose costs [30]. This is possible, and though we did not evaluate R* or fitness at multiple evolutionary end-points, fitness may continue increasing under directional selection for tens of thousands of generations [59]. However, if trade-offs do not emerge within 285 generations of low-resource selection, natural populations of phytoplankton evolving in response to seasonal or annual variation in nutrient availability may not be expected to be optimizing along trade-off axes in competitive ability for different resources. Finally, mutations affecting any particular resource requirement may generally be more likely to be synergistically pleiotropic than neutral or antagonistic. Given the degree of metabolic interrelatedness of resource acquisition and metabolic pathways in phytoplankton, this is plausible and deserves further investigation.

Patterns of genotypic variation across populations revealed negative correlations between fitness at low and high resource supply for a given resource (gleaner-opportunist trade-offs, Figure 1A), and positive correlations between competitive abilities for different resources (Figure 3, ESM Tables 4 - 6), suggesting that the evolution of competitive ability could be constrained by genetic correlations between multiple resource traits under selection. The genetic correlations between different resource traits could explain the positively associated trait changes (i.e. improvements in multiple minimum resources requirements simultaneously). In addition, unmeasured traits could be involved in the trade-off, resulting in a positive genetic covariance between any two resource traits [18]. When testing for trade-offs, we accounted for concurrent variation in cell size, but other fitness-correlated traits, such as resistance to grazers or pathogens [60,61], may be involved in the trade-offs.

Traits relevant to competitive ability, such as cell size, are known to change as a result of phenotypic plasticity and evolutionary adaptation [62]. We have demonstrated that adaptation in response to resource limitation and salt stress can alter competitive traits sufficiently to change the predicted outcome of competition. Adaptation to different environments caused competitive traits to diverge and enable coexistence. Contrary to our expectations, we found that coexistence was equally likely among two populations selected in different environments as two populations selected in the same environment. This may be explained by the fact that even small differences in the magnitude of adaptive trait change in the same environment can be sufficient to enable predicted coexistence (i.e. under P-limitation, P^*^ for one competitor decreases slightly more than the P^*^ for the other competitor). The changes in resource ratios and salt levels represented in our different selection environments are on the same order of magnitude as gradients of resource ratios and salinity in natural environments [63]. This means that predictions of the outcomes of competition should incorporate the potential for evolutionary changes to influence competitive dynamics [16].

Our results are directly relevant to understanding eco-evolutionary feedbacks in competitively structured communities. Theory predicts that species converge in their resource-use traits when competing for essential resources [15,16]. This expectation, however, depends on two critical assumptions. These assumptions are that species’ consumption vectors remain fixed and that competitive abilities for different limiting resources trade off. While we did not grow pairs of populations together in the evolution experiment, the resource limitation treatments mimicked the effects of a better competitor for the limiting resource, while avoiding exclusion of the weaker competitor. Our results do not provide empirical support for either of the assumptions above, suggesting that theoretical predictions of evolutionary adaptation under essential resource competition may need to be revised [15,16].

Understanding patterns of biodiversity and coexistence requires accounting for past and current evolutionary changes in species’ competitive traits. While macro-evolutionary patterns show trade-offs in species’ resource-use traits, we found that positively correlated adaptive trait changes drive within-species responses to resource limitation, altering the expected outcome of competition. Such micro-evolutionary changes in species’ competitive abilities should to be considered if we are to improve our predictions of competitive interactions and community dynamics in a changing world.

## Acknowledgments

We thank the following people who helped us maintain cultures, and collect and process the data: C. Carvalho, G. Siegrist, P. Ganesanandamoorthy, L. Rihakova, D. Steiner, E. Burmeister, M. Thali and J-C. Walser. Thanks to J. Jokela for helpful comments on an earlier version of the manuscript. The genomic data produced and analyzed in this paper were generated in collaboration with the Genetic Diversity Centre (GDC), ETH Zurich. We thank J. Martiny and two anonymous reviewers for constructive comments and suggestions.

## Data and code accessibility

Data and code are available at https://github.com/JoeyBernhardt/chlamee-r-star and will be deposited in the Dryad digital data repository should the manuscript be accepted. DNA sequences have been deposited in the Sequence Read Archive (SRA) under the BioProject ID PRJNA558172.

## References

1. Tilman D. 1982 Resource Competition and Community Structure. Princeton, New Jersey: Princeton University Press.

2. Edwards KF, Litchman E, Klausmeier CA. 2013 Functional traits explain phytoplankton responses to environmental gradients across lakes of the United States. Ecology 94, 1626–1635.

3. Keddy PA. 2015 Competition. eLS. (doi:10.1002/9780470015902.a0003162.pub2)

4. Chesson P. 2000 Mechanisms of maintenance of species diversity. Annu. Rev. Ecol. Syst. 31, 343–366.

5. Miller et al. 2005 A Critical Review of Twenty Years’ Use of the Resource-Ratio Theory. Am. Nat. 165, 439–448.

6. Hart SP, Turcotte MM, Levine JM. 2019 Effects of rapid evolution on species coexistence. Proc. Natl. Acad. Sci. U. S. A. 116, 2112–2117.

7. Lankau RA. 2011 Rapid Evolutionary Change and the Coexistence of Species. Annu. Rev. Ecol. Evol. Syst. 42, 335–354.

8. Germain RM, Williams JL, Schluter D, Angert AL. 2018 Moving Character Displacement beyond Characters Using Contemporary Coexistence Theory. Trends in Ecology & Evolution. 33, 74–84. (doi:10.1016/j.tree.2017.11.002)

9. Ingram T, Svanbäck R, Kraft NJB, Kratina P, Southcott L, Schluter D. 2012 Intraguild predation drives evolutionary niche shift in threespine stickleback. Evolution 66, 1819–1832.

10. Pfennig DW, Pfennig KS. 2010 Character Displacement and the Origins of Diversity. The American Naturalist. 176, S26–S44. (doi:10.1086/657056)

11. Dayan T, Simberloff D. 2005 Ecological and community-wide character displacement: the next generation. Ecol. Lett. 8, 875–894.

12. Schluter D. 2000 The ecology of adaptive radiation. OUP Oxford.

13. Grant PR, Grant BR. 2006 Evolution of character displacement in Darwin’s finches. Science 313, 224–226.

14. MacArthur R, Levins R. 1967 The limiting similarity, convergence, and divergence of coexisting species. Am. Nat. 101, 377–385.

15. Abrams PA. 1987 Alternative models of character displacement and niche shift. I. Adaptive shifts in resource use when there is competition for nutritionally nonsubstitutable resources. Evolution 41, 651–661.

16. Fox JW, Vasseur DA. 2008 Character convergence under competition for nutritionally essential resources. Am. Nat. 172, 667–680.

17. Lande R, Arnold SJ. 1983 The measurement of selection on correlated characters. Evolution 37, 1210–1226.

18. Blows MW, Hoffmann AA. 2005 A reassessment of genetic limits to evolutionary change. Ecology

19. Litchman E, Klausmeier CA, Schofield OM, Falkowski PG. 2007 The role of functional traits and trade-offs in structuring phytoplankton communities: scaling from cellular to ecosystem level. Ecol. Lett. 10, 1170–1181.

20. Edwards KF, Klausmeier CA, Litchman E. 2011 Evidence for a three-way trade-off between nitrogen and phosphorus competitive abilities and cell size in phytoplankton. Ecology 92, 2085–2095.

21. Litchman E, Klausmeier CA. 2008 Trait-Based Community Ecology of Phytoplankton. Annu. Rev. Ecol. Evol. Syst. 39, 615–639.

22. Grover JP, Hudziak J, Grover JD. 1997 Resource Competition. Springer Science & Business Media.

23. Kirk KL. 2002 Competition in variable environments: experiments with planktonic rotifers. Freshwater Biology. 47, 1089–1096. (doi:10.1046/j.1365-2427.2002.00841.x)

24. Strzepek RF, Harrison PJ. 2004 Photosynthetic architecture differs in coastal and oceanic diatoms. Nature 431, 689–692.

25. Rhee G--U, Gotham IJ. 1981 The effect of environmental factors on phytoplankton growth: Light and the interactions of light with nitrate limitation 1. Limnol. Oceanogr. 26, 649–659.

26. Grover JP. 1997 Resource Competition. (doi:10.1007/978-1-4615-6397-6)

27. Monod J. 1949 The growth of bacterial cultures. Annu. Rev. Microbiol. 3, 371–394.

28. Bono LM, Smith LB Jr, Pfennig DW, Burch CL. 2017 The emergence of performance trade-offs during local adaptation: insights from experimental evolution. Mol. Ecol. 26, 1720–1733.

29. Shoval O, Sheftel H, Shinar G, Hart Y, Ramote O, Mayo A, Dekel E, Kavanagh K, Alon U. 2012 Evolutionary trade-offs, Pareto optimality, and the geometry of phenotype space. Science 336, 1157–1160.

30. Li Y, Petrov DA, Sherlock G. 2019 Single nucleotide mapping of trait space reveals Pareto fronts that constrain adaptation. Nat Ecol Evol 3, 1539–1551.

31. Tamminen M, Betz A, Pereira AL, Thali M, Matthews B, -F. Suter MJ, Narwani A. 2018 Proteome evolution under non-substitutable resource limitation. Nature Communications. 9. (doi:10.1038/s41467-018-07106-z)

32. Geider R, La Roche J. 2002 Redfield revisited: variability of C:N:P in marine microalgae and its biochemical basis. Eur. J. Phycol. 37, 1–17.

33. Sterner RW, Elser JJ. 2002 Ecological stoichiometry: the biology of elements from molecules to the biosphere. Princeton University Press.

34. Goho S, Bell G. 2000 The ecology and genetics of fitness in Chlamydomonas. IX. The rate of accumulation of variation of fitness under selection. Evolution 54, 416–424.

35. McGee LW, Sackman AM, Morrison AJ, Pierce J, Anisman J, Rokyta DR. 2016 Synergistic Pleiotropy Overrides the Costs of Complexity in Viral Adaptation. Genetics 202, 285–295.

36. Frachon L et al. 2017 Intermediate degrees of synergistic pleiotropy drive adaptive evolution in ecological time. Nat Ecol Evol 1, 1551–1561.

37. Bell G. 2008 Selection: the mechanism of evolution. Oxford University Press on Demand.

38. Hughes AR, Inouye BD, Johnson MT, Underwood N, Vellend M. 2008 Ecological consequences of genetic diversity. Ecol. Lett. 11, 609–623.

39. Barrett RDH, Schluter D. 2008 Adaptation from standing genetic variation. Trends Ecol. Evol. 23, 38–44.

40. Bell G, Collins S. 2008 Adaptation, extinction and global change. Evol. Appl. 1, 3–16.

41. Goho S, Bell G. 2000 Mild environmental stress elicits mutations affecting fitness in Chlamydomonas. Proc. R. Soc. Lond. B Biol. Sci. 267, 123–129.

42. Velicer GJ, Lenski RE. 1999 Evolutionary trade-offs under conditions of resource abundance and scarcity: experiments with bacteria. Ecology 80, 1168–1179.

43. Velicer GJ. 1999 Pleiotropic effects of adaptation to a single carbon source for growth on alternative substrates. Appl. Environ. Microbiol. 65, 264–269.

44. Lahti DC, Johnson NA, Ajie BC, Otto SP, Hendry AP, Blumstein DT, Coss RG, Donohue K, Foster SA. 2009 Relaxed selection in the wild. Trends Ecol. Evol. 24, 487–496.

45. Kilham SS, Kreeger DA, Lynn SG, Goulden CE, Herrera L. 1998 COMBO: a defined freshwater culture medium for algae and zooplankton. Hydrobiologia 377, 147–159.

46. Huisman J, Weissing FJ. 1994 Light-limited growth and competition for light in well-mixed aquatic environments: an elementary model. Ecology 75, 507–520.

47. Elzhov TV, Mullen KM, Spiess A-N, Bolker B. 2013 minpack.lm: R interface to the Levenberg-Marquardt nonlinear least-squares algrothim found in MINPACK, pluss support for bounds. See https://cran.r-project.org/web/packages/minpack.lm/minpack.lm.pdf.

48. Eilers PHC, Peeters JCH. 1988 A model for the relationship between light intensity and the rate of photosynthesis in phytoplankton. Ecol. Modell. 42, 199–215.

49. Baty F, Ritz C, Charles S, Brutsche M, Flandrois J-P, Delignette-Muller M-L. 2015 A Toolbox for Nonlinear Regression in R : The Package nlstools. J. Stat. Softw. 66, 1–21.

50. Oksanen J et al. 2013 Package ‘vegan’. Community ecology package, version 2.

51. R Core Team. 2019 R: A Language and Environment for Statistical Computing.

52. Klausmeier CA, Litchman E, Daufresne T, Levin SA. 2004 Optimal nitrogen-to-phosphorus stoichiometry of phytoplankton. Nature 429, 171–174.

53. Gurevitch J, Morrow LL, Wallace A, Walsh JS. 1992 A Meta-Analysis of Competition in Field Experiments. Am. Nat. 140, 539–572.

54. Adler PB, Hillerislambers J, Levine JM. 2007 A niche for neutrality. Ecol. Lett. 10, 95–104.

55. Kronholm I, Bassett 1. Andrew, Baulcombe D, Collins S. In press. Epigenetic and Genetic Contributions to Adaptation in. Evolution 34, 2285–2306.

56. Schluter D. 1996 ADAPTIVE RADIATION ALONG GENETIC LINES OF LEAST RESISTANCE. Evolution 50, 1766–1774.

57. Guo J et al. 2018 Specialized proteomic responses and an ancient photoprotection mechanism sustain marine green algal growth during phosphate limitation. Nat Microbiol 3, 781–790.

58. Litchman E, de Tezanos Pinto P, Klausmeier CA, Thomas MK, Yoshiyama K. 2010 Linking traits to species diversity and community structure in phytoplankton. In Fifty years after the ‘“Homage to Santa Rosalia”‘: Old and new paradigms on biodiversity in aquatic ecosystems (eds L Naselli-Flores, G Rossetti), pp. 15–28. Dordrecht: Springer Netherlands.

59. Lenski RE et al. 2015 Sustained fitness gains and variability in fitness trajectories in the long-term evolution experiment with Escherichia coli. Proceedings of the Royal Society B: Biological Sciences 282, 20152292.

60. Yoshida T, Hairston NG Jr, Ellner SP. 2004 Evolutionary trade-off between defence against grazing and competitive ability in a simple unicellular alga, Chlorella vulgaris. Proceedings of the Royal Society B - Biological Sciences 271, 1947–1953.

61. Bowers Roger G., Boots Michael, Begon Michael. 1994 Life-history trade-offs and the evolution of pathogen resistance: competition between host strains. Proceedings of the Royal Society of London. Series B: Biological Sciences 257, 247–253.

62. Malerba ME, Marshall DJ. 2019 Size-abundance rules? Evolution changes scaling relationships between size, metabolism and demography. Ecology Letters. 22, 1407–1416. (doi:10.1111/ele.13326)

63. Wetzel RG. 2001 Limnology: Lake and River Ecosystems. Gulf Professional Publishing.

