## Appendix A: Supplementary Methods for "The evolution of competitive ability for essential resources"

### Electronic Supplementary Materials: Appendix A

### Supplementary methods

*Evolution experiment*

We obtained a strain of *C. reinhardtii* (CC1690 wild type mt+) from the Chlamydomonas Center (chlamycollection.org). We then grew this strain in semi-continuous liquid batch culture on COMBO freshwater medium [1], without vitamins, silica and animal trace elements, which are unnecessary for growing green algae. We maintained batch cultures for several months before the start of the evolution experiment. We then plated the cultures onto agar. From the agar plates we selected four random colonies, derived from single cells and inoculated them into liquid COMBO freshwater medium (hereafter referred to as Anc 2, Anc 3, Anc 4 and Anc 5). We then inoculated the chemostats (28 mL total volume) with one of the four monoclonal populations or a genetically diverse population of the original CC1690 population. All populations were composed of a single mating type (+), precluding the possiblity of sex during the experiment. Here we use the term 'population' to refer to each of Anc 2, Anc 3, Anc 4, Anc 5, cc1690 ('ancestors') and their descendant populations ('descendants'), which are the populations evolved in one of seven experimental environments. In total there were five ancestral populations, and 32 descendant populations (three were lost to contamination).

We randomly assigned one chemostat of each of the 5 ancestral populations (Anc 2-5 and CC1690) to one of seven treatments: COMBO, (which we call C, containing 1000 uM N and 50 uM P), nitrogen limitation (N, 10 uM N and 50 uM P), phosphorus limitation (P, 1000 uM N, 0.5 uM P), light limitation (L, 5 µmol photons $m^{-2}s^{-1}$of light,1000 uM N and 50 uM P), salt stress (S, 8g/L NaCl, 1000 uM N and 50 uM P), biotically-depleted medium (i.e. medium previously used to grow seven other species of phytoplankton, (B)), and a combination of salt stress and biotically-depleted medium [2] (BS, 8g/L NaCl). We prepared the biotically-depleted medium (B) by growing seven other species of freshwater algae (*Cosmarium botrytis, Kirchneriella subcapitata, Pediastrum boryanum, Pediastrum duplex, Scenedesmus acuminatus, Staurastrum punctulatum, Tetraedron minimum*) individually in 4L batch culture in replete COMBO (Kilham et al. 1998). We then removed the phytoplankton biomass from each batch culture via filtration, sterilized the filtrate by autoclaving, and mixed the filtrate from each of the seven cultures. The mixture of filtrates is what we called the 'biotically depleted medium' (B). We stored the biotically-depleted medium in the cold (4°C) and dark until use during the experiment.

Each chemostat received daily sterile media replacement at a dilution rate of 56% per day via an automated peristaltic pump and was continuously mixed and aerated to prevent heterogeneity in resource availability. Chemostats each contained a magnetic stir bar at their base and were stirred continuously using a stir plate. The combination of continuous sterile air inflow and stirring kept the chemostats well mixed and minimized wall growth, and in this way minimized spatial heterogeneity. We maintained chemostats at 20°C and illuminated under 90 µmol photons $m^{-2}s^{-1}$of light (except the light-limitation treatment) on an 18h light: 6h dark cycle.

The biotically-depleted medium was used to investigate the influence that a biodiverse community may have on the evolution of a species by simultaneously depleting the availability of multiple dissolved resources. The salt stress treatment consisted of increasing concentrations of NaCl. We used salt-stress as a point of reference by which to compare the influence of a resource limitation. Salt stress is known to induce evolutionary adaptation (i.e. greater salt tolerance) in *C. reinhardtii [3],* and in this way we could compare adaptation to limiting resources to another type of stress. We maintained the 'C' cultures in full COMBO medium for the duration of the experiment. Resource-limitation and salt-stress increased incrementally each month until a final, highly-stressful concentration was achieved (ESM Figure S3). The experiment ran for 285 days (~285 generations), after which the evolved populations were harvested and plated onto agar.

***Culture conditions and acclimation***

Prior to $R^*$ assays, we transferred populations from agar plates where they had been maintained for long-term storage after the selection experiment under low light (amount) and temperature (12°C) to limit growth to COMBO medium [1] and grew them at 20°C and 140 µmol light (hereafter "standard conditions") on a 24 hour light cycle for three days until they reached mid exponential phase. Before the start of each $R^*$ experiment, we allowed each population to acclimate to a relatively low and high resource level for three days. The low resource acclimation conditions were set to match the lowest resource level in the $R^*$ experiments so as to minimize transfer of nutrients from the nutrient replete culture media to the experimental populations. The high resource acclimation level was set halfway between the lowest and highest resource level in the $R^*$ experiments. The low resource acclimation levels were for light, nitrogen and phosphorus: 8μmols/m$^2$/s, 5 uM N and 0.5 uM P. The high resource acclimation levels were for light, nitrogen and phosphorus: 42 μmols/m$^2$/s, 80 uM N, 8 uM P.

***Competitive trait assays***

We diluted the 'low' and 'high' resource acclimation cultures to low cell concentrations in 50 mL falcon tubes with COMBO media containing N and P at one of ten resource levels. After diluting each population to very low density (measured as 10 chlorophyll-a relative fluorescence units ("RFU")) at each resource level, we transferred the cultures to the inner 60 wells of 96 well plates (n = 4 replicates per population per resource level, 125 uL per well), covered the plates with a Breathe-Easy sealing membrane (Sigma-Aldrich), and moved them to 20°C temperature-controlled incubators (Multitron, Infors HT, Switzerland), which we set to rotate at 100 rpm. We filled outer wells with COMBO to prevent evaporative losses across the plate. We then tracked their growth by measuring chlorophyll-a fluorescence in RFU (excitation=435 nm and emission=685 nm) over time using a Biotek Cytation 5 plate-reader. We used chlorophyll-a fluorescence because it can be used as a proxy for algal biomass [4], particularly during exponential growth from low density. We measured RFUs two or three times a day for three days, long enough to capture the exponential growth phase at all resource levels. For the $N^*$ and $P^*$ experiments, cultures were illuminated at 140.6 μmol photons m-2·s-1 of photosynthetically active radiation ('PAR'); for the $I^*$ experiments light levels were as described below.

For the nitrogen $R^*$ experiment, the N levels were: 5, 10, 20, 40, 60, 80, 100, 400, 600 and 1000 um N. For the phosphorus $R^*$ experiment, the P levels were 0.5, 1, 2, 4, 6, 8, 10, 20, 35 and 50 uM P. For the light $R^*$ experiment, N and P were 1000 uM N and 50 uM P respectively, and light was supplied at one of ten levels: 0.25, 1.5, 5, 12.5, 27.5, 50, 82.5, 125, 175 and 250 μmol photons m$^{-2}$·s$^{-1}$ of PAR. We manipulated light levels in the light experiment using neutral density filters (Solar Graphics™, Clearwater, Florida), which alter the total amount of light supplied without changing light spectrum. We mounted the light filters onto opaque frames, which fit over the plates and prevented unmeasured light from entering the wells from the sides of the plates. We measured experimental light intensities under the filters using a Skye PAR Quantum sensor.

***Salt tolerance assays***

Similar to the methods for the $R^*$ assays, all ancestral and descendant populations were first transferred from storage on agar plates to liquid batch cultures and grown in standard conditions. They were then transferred to liquid culture to start an acclimation phase in which each of the populations was subjected to one of five levels of NaCl: 0, 2, 4, 6, and 8 g/L for four days. Each of the populations was then diluted to achieve a final inoculation density of 50 RFU. Populations from each of the acclimation levels were used to inoculate assay cultures with the same level of salt, or 1 g/L more (i.e. 0 was used to inoculate 0 and 1 g/L, 2 to inoculate 2 and 3 g/L, etc.). For the final growth rate assays, each population was grown in 10 mL of medium in 6-well plates, with a single replicate per population x salt level. We estimated salt tolerance by growing populations over a salt gradient of 0, 1, 2, 3, 4, 5, 6, 7, 8, 9, 10 g/L for nine days. To estimate growth rates, RFUs were measured once per day for nine days.

*Estimating consumption vectors via stoichiometry*

We quantified the consumption vectors for nitrogen and phosphorus for each population. The consumption vector [5] quantifies the amount of each of these two resources used by each individual consumes per unit time. We estimated these vectors by measuring the ratio of phosphorus to nitrogen in the biomass of each population as it was growing exponentially [6]. Stoichiometry during the exponential phase primarily reflects the structural pool of nutrients (vs. the storage pool) [7]. We started by inoculating each population into a 400 mL tissue culture flask with COMBO [1]. We then allowed these populations to grow for approximately 1.5 days under standard conditions until they reached their mid-exponential phase. We then harvested the algal biomass by filtering each culture onto one ashed (400 °C) and pre-massed Whatman ® glass microfiber filter (grade GF/F 47 mm) and one 25mm Whatman glass microfiber filter. We then dried the filters dried in an oven overnight at 60 °C, and post-massed the 47mm filters to obtain an estimate of total dry biomass per mL of culture filtered. The 47mm filter was used to estimate the elemental carbon and nitrogen content of the biomass on an Elementar vario PYRO cube EA-IRMS, and the 25 mm filter was used to estimate phosphorus content using Skalar San++Continuous Flow P/N analyser. The phosphorus samples were first digested and completely oxidized using a peroxydisulfate solution. We diluted the digested samples 1:20 before being run on the P/N analyser.

***Estimating cell size***

After the final RFU measurements, we fixed the populations in each well by adding a 10% glutaraldehyde fixative (get details) solution, and stored the plates at 4°C until later analysis. To estimate cell size, we took Brightfield photos of the base of each well at 10x on a BioTek Cytation 5 imaging plate reader, from which we extracted cell length (using Gen5 software (BioTek version 2.0), which we converted to biovolume, assuming the cells were spheres (i.e. $4/3 \times \pi \times radius^3$).

***Quantifying genetic changes associated with selection environments***

To gain insight into the genetic responses of *C. reinhardtii* to the selection environments, we prepared Illumina HiSeq libraries of the ancestors and descendants. The ancestral populations of all 4 clonal populations and the original cc1690 population were plated onto Sueoka's high salt agar [8] and grown on agar plates for 1 month. We harvested the lawn of cells by scraping the agar and placing the biomass into microfuge tubes before performing the DNA extraction. The descendant populations were grown in 50 mL liquid batch cultures in COMBO medium in standard conditions for one week. Due to low level bacterial contamination, and to ensure that sequences were highly enriched by *C. reinhardtii*, these cultures were subjected to an antibiotic treatment of 50 mg/L ampicillin and 50 mg/L tetracycline overnight (<24 hours), before harvesting the cells for DNA extraction [9]. The cells were then harvested by centrifugation at 4,000 rpm.  The DNA extraction protocol was adapted from the Plant Lab protocol (Institute of Life Sciences, Scuola Superiore Sant'Anna, Pisa Italy and [10]).

DNA sequencing libraries were prepared using the Bioo Scientific NEXTflex Rapid Illumina DNA-Seq Library Prep Kit according to the standard protocol.  DNA sequencing was performed on Illumina HiSeq 2500 version 4 using 125 bp paired ends (250 sequencing cycles). After sequencing the read quality was verified using FastQC version 0.11.2. Adaptor and PhiX cleaning were performed using BBDuk version 35.43, using k-mer size 20 for the former. Quality filtering was performed using PRINSEQ version 0.20.4 with a minimum read length of 50 bp, GC range of 15-85% and minimum mean quality score 5. The quality-filtered reads were aligned against the *C. reinhardtii* reference version 5.0 [11] using Bowtie2 version 2.2.5. Variants were called from the resulting BAM files using Freebayes version 1.1.0. The resulting VCF files were quality-filtered using bcftools version 1.4 to select above SNP quality 20 and excluding any SNPs closer to 10 bp from any INDEL due to known read mapping errors around such mutations. The filtered VCF files further processed in R version 3.5.1 using library Tidyverse version 1.2.1 and Bioconductor package VariantAnnotation version 1.28.11. The mutations between the ancestors and descendants were determined by comparing their SNP profiles, determined by comparison to the *C. reinhardtii* cc503 mt+, reference version 5.0, using custom R scripts. The R code for sequence processing is available at https://github.com/joeybernhardt/chlamee-r-star/blob/master/genomics/workflow.R. The DNA sequences have been deposited in the Sequence Read Archive (SRA) under the BioProject ID PRJNA558172.

### References


1. Kilham SS, Kreeger DA, Lynn SG, Goulden CE, Herrera L. 1998 COMBO: a defined freshwater culture medium for algae and zooplankton. *Hydrobiologia* **377**, 147–159.

2. Tamminen M, Betz A, Pereira AL, Thali M, Matthews B, -F. Suter MJ, Narwani A. 2018 Proteome evolution under non-substitutable resource limitation. *Nat. Commun.* **9**, 4650. (doi:10.1038/s41467-018-07106-z)

3. Lachapelle J, Bell G, Colegrave N. 2015 Experimental adaptation to marine conditions by a freshwater alga. *Evolution* **69**, 2662–2675. (doi:10.1111/evo.12760)

4. Clesceri LS. 1998 *Standard Methods for the Examination of Water and Wastewater*. American Public Health Association. See https://play.google.com/store/books/details?id=pv5PAQAAIAAJ.

5. Tilman D. 1982 Resource competition and community structure. *Monogr. Popul. Biol.* **17**, 1–296.

6. Keddy PA. 2015 Competition. *eLS*. (doi:10.1002/9780470015902.a0003162.pub2)

7. Geider R, La Roche J. 2002 Redfield revisited: variability of C:N:P in marine microalgae and its biochemical basis. *Eur. J. Phycol.* **37**, 1–17. (doi:10.1017/S0967026201003456)

8. Sueoka N. 1960 Mitotic replication of deoxyribonucleic acid in Chlamydomonas reinhardtii. *Proc. Natl. Acad. Sci. U. S. A.* **46**, 83–91. (doi:10.1073/pnas.46.1.83)

9. Andersen RA. 2005 *Algal Culturing Techniques*. Academic Press. See https://play.google.com/store/books/details?id=9NADUHyFZaEC.

10. Ness RW, Morgan AD, Colegrave N, Keightley PD. 2012 Estimate of the spontaneous mutation rate in Chlamydomonas reinhardtii. *Genetics* **192**, 1447–1454. (doi:10.1534/genetics.112.145078)

11. Merchant SS *et al.* 2007 The Chlamydomonas genome reveals the evolution of key animal and plant functions. *Science* **318**, 245–250. (doi:10.1126/science.1143609)
