## Appendix B: Supplementary Figures for "The evolution of competitive ability for essential resources"

### Electronic Supplementary Materials: Appendix B

#### Supplementary Figures
#### Figures S1 - S17

### The evolution of competitive ability for essential resources

Joey R. Bernhardt[1*], Pavel Kratina[2], Aaron Pereira[1], Manu Tamminen[3], Mridul K. Thomas4, Anita Narwani[1]

[1]Aquatic Ecology Department, Eawag, Überlandstrasse 133, CH-8600 Dübendorf, Switzerland

[2]School of Biological and Chemical Sciences, Queen Mary University of London, Mile End Road, London E1 4NS, United Kingdom.

[3]Department of Biology, University of Turku, Natura, University Hill, 20014 Turku, Finland

[4]Centre for Ocean Life, DTU Aqua, Technical University of Denmark, Kongens Lyngby, Denmark

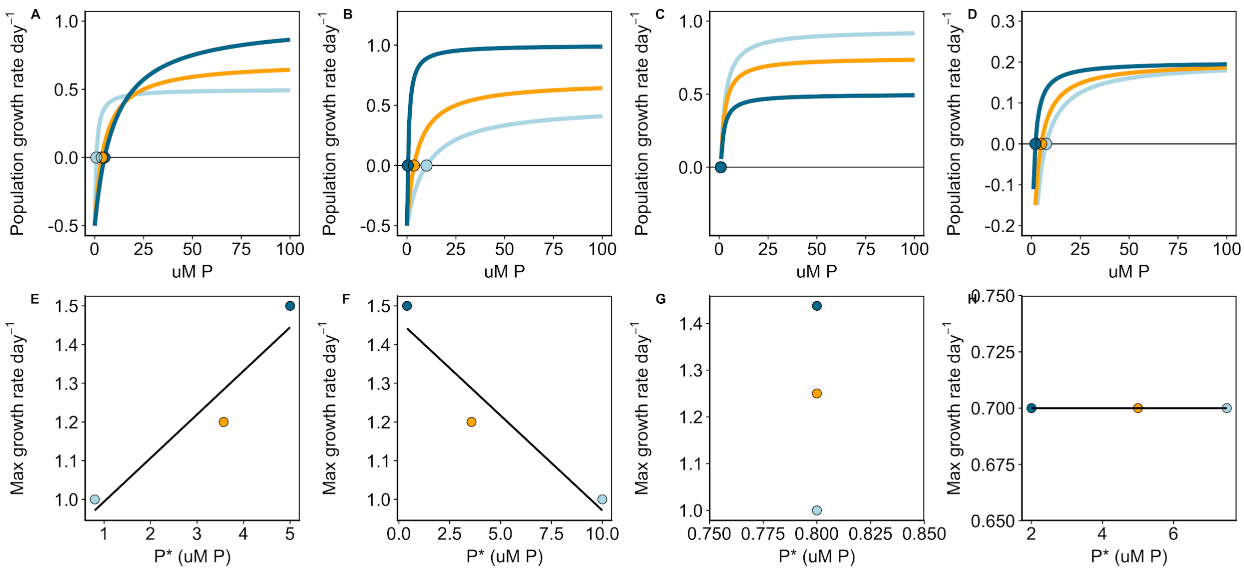


**Figure S1.** Resource-dependent growth can be characterized by a Monod curve [1]. Monod curves characterizing possible the relationships between population growth rates and resource concentration (A-D). Variation among populations (illustrated by different colours) can be characterized by: a gleaner-opportunist trade-off scenario [2] (A and E), a fast-slow continuum scenario [3,4], or vertical shift) (B and F), vertical shift associated with no variation in minimum resource requirement (C, G) or variation in minimum resource requirement with no associated variation in maximum growth rate. Lastly, variation in the Monod curve may show constraint or correlation among parameters (i.e. all shapes are possible). Genetic or biophysical constraints on how the parameters of the Monod curves vary constrain evolutionary outcomes.

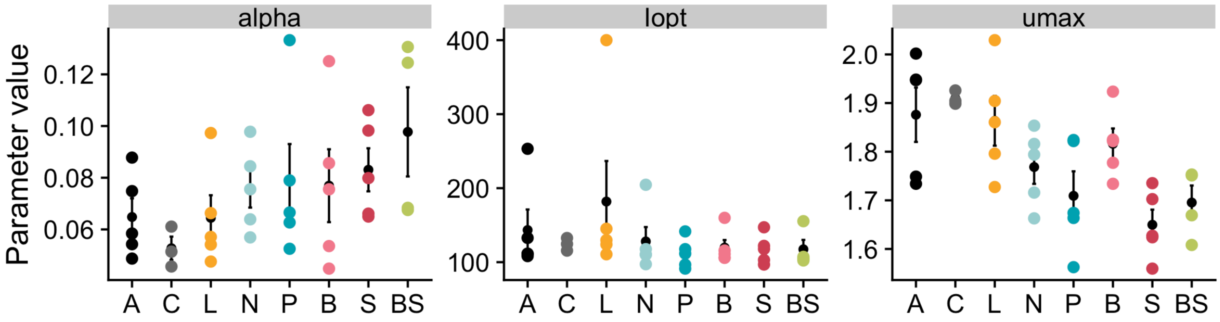


**Figure S2.** Parameter estimates from an Eilers-Peeters [5] curve fit to population growth rate over a light gradient. In the Eilers and Peeters dynamic model of photoinhibition of photosynthesis: , where is the growth rate (day-1) and a function of photon flux density (I, photons m2s-1), is the maximum growth rate at Iopt, and is the initial slope of the curve. A: ancestors, C: COMBO, L: light-limited, P: P-limited, N: N-limited, B: biotically depleted media, S: high salt, BS: biotically depleted and high salt.

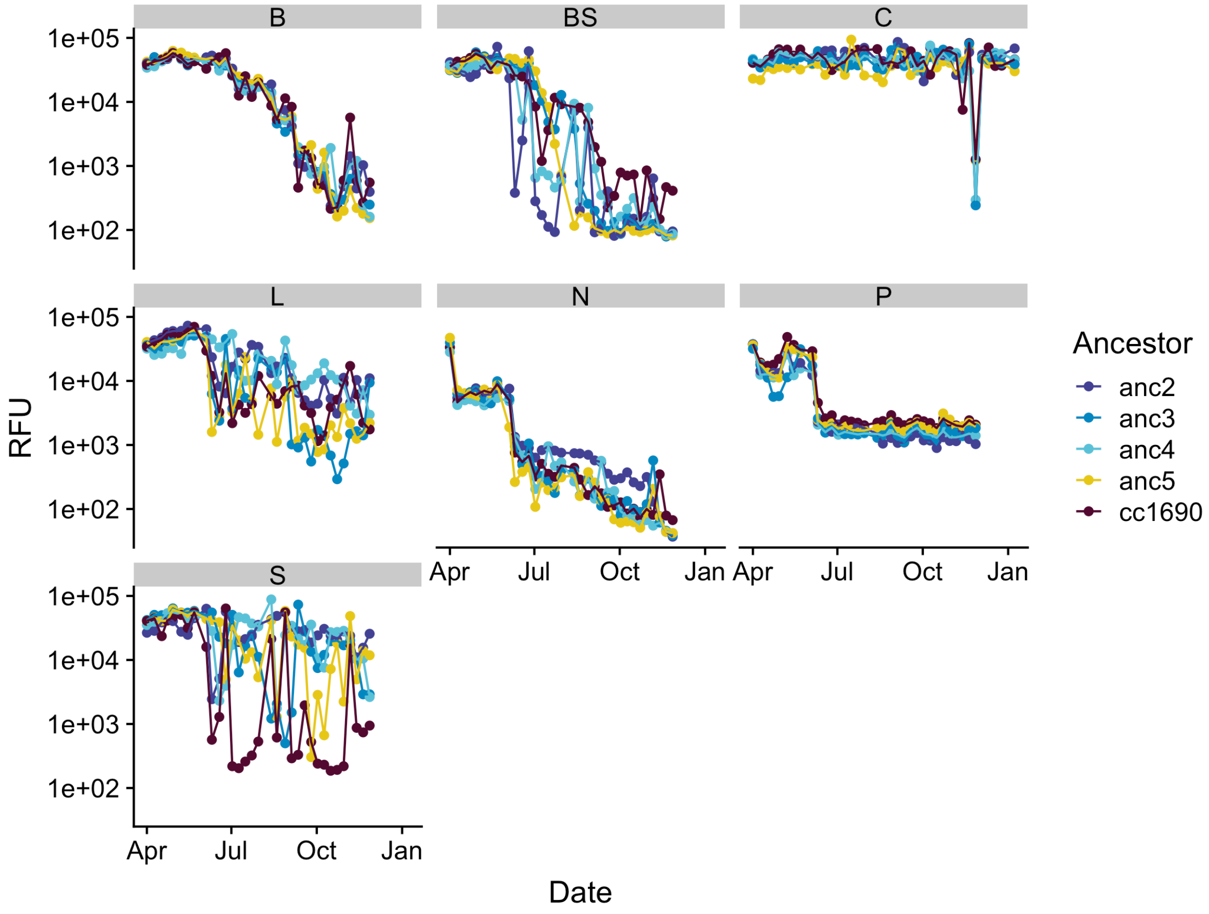


**Figure S3.** Population abundance trajectories over the evolution experiment estimated via Relative Fluorescence Units (RFU). Note that the y-axis is on a log scale. Panels correspond to selection environments (B: biotically depleted, BS: Biotically-depleted + high salt, C: COMBO, L: light limitation, N: nitrogen limitation, P: phosphorus limitation, S: high salt).

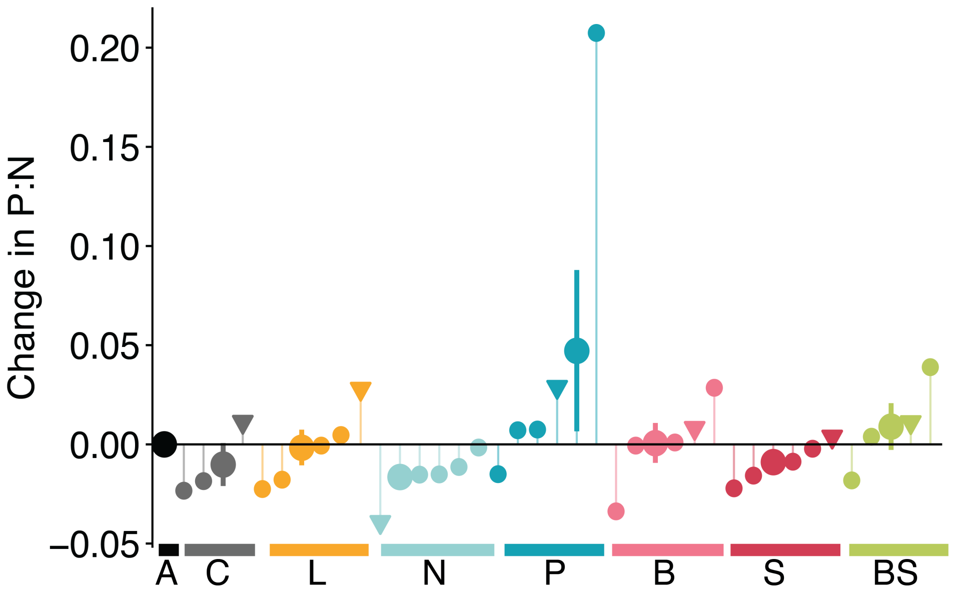


**Figure S4.** Change in consumption vectors, estimated as P:N molar ratio of the descendant populations relative to their ancestors. A: ancestors, C: COMBO, L: light-limited, P: P-limited, N: N-limited, B: biotically depleted media, S: high salt, BS: biotically depleted and high salt. Small points correspond to individual populations and large points correspond to the average change of all populations in a given environment. Populations represented with a triangle are the genotypically diverse populations (cc1690). Error bars correspond to SE.

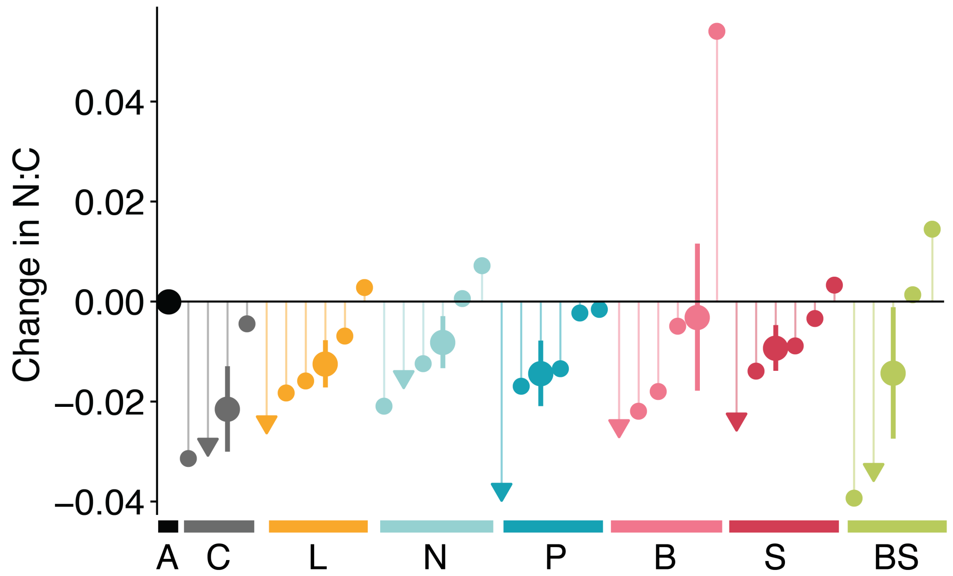


**Figure S5.** Change in population biomass N:C of the descendants relative to their ancestors. C: COMBO, L: light-limited, P: P-limited, N: N-limited, B: biotically depleted media, S: high salt, BS: biotically depleted and high salt. Small points correspond to individual populations and large purple points correspond to the average change of all populations in a given environment. Error bars correspond to SE.

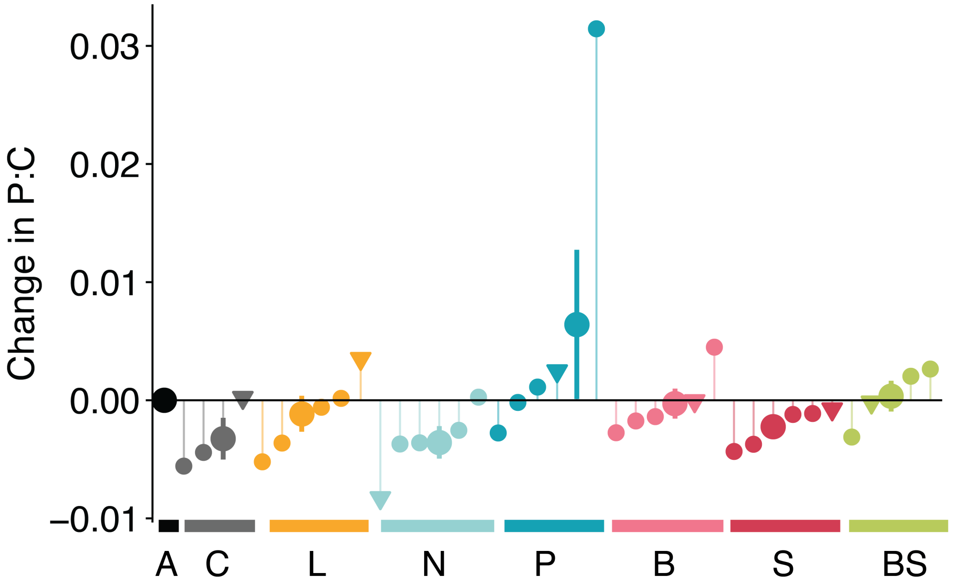


**Figure S6.** Change in population biomass P:C of the descendants relative to their ancestors. C: COMBO, L: light-limited, P: P-limited, N: N-limited, B: biotically depleted media, S: high salt, BS: biotically depleted and high salt. Small points correspond to individual populations and large purple points correspond to the average change of all populations in a given environment. Error bars correspond to SE.

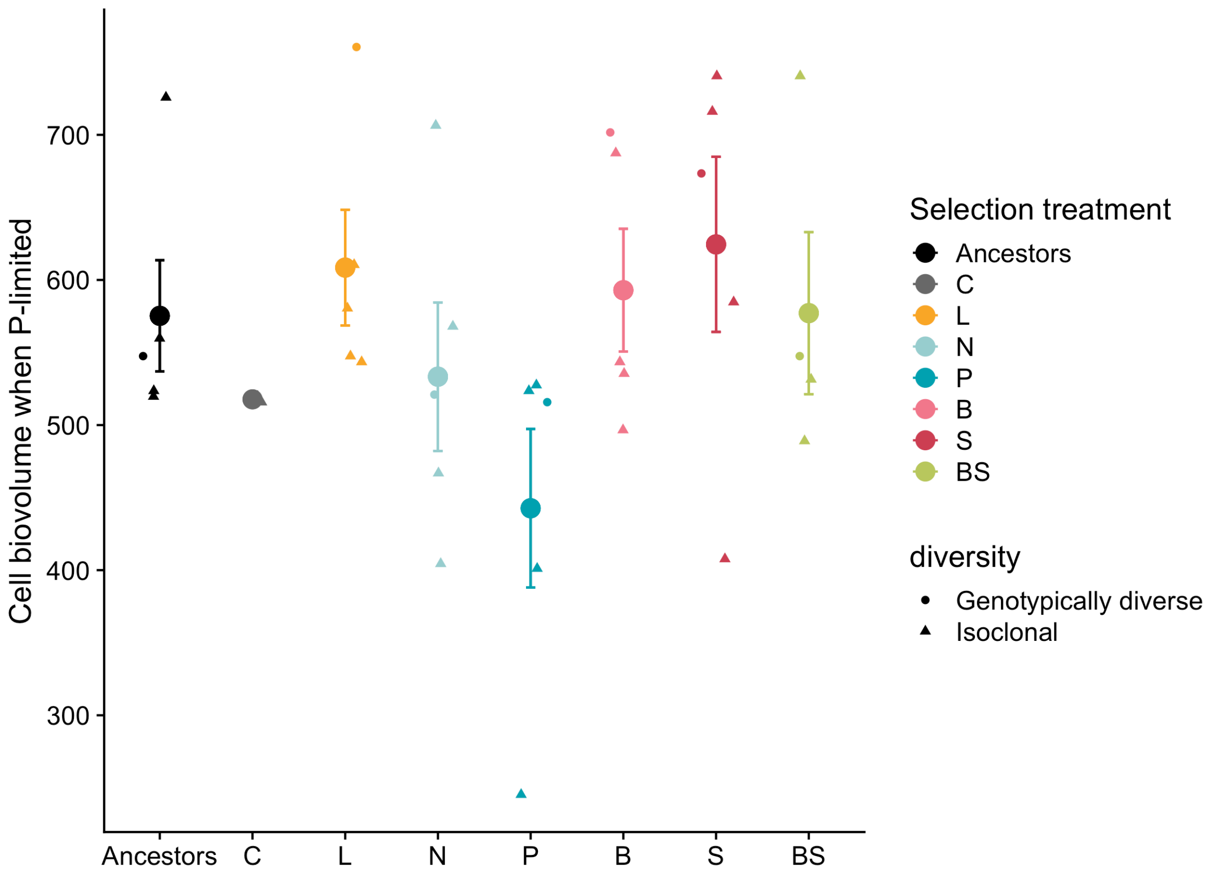


**Figure S7.** Cell biovolume (um$^3$) estimated in common garden conditions in batch culture in 0.5 uM P media. Large points and error bars correspond to the average and standard error across replicate populations. Colours correspond to the selection environment: A: ancestors, C: COMBO, L: light-limited, P: P-limited, N: N-limited, B: biotically depleted media, S: high salt, BS: biotically depleted and high salt.

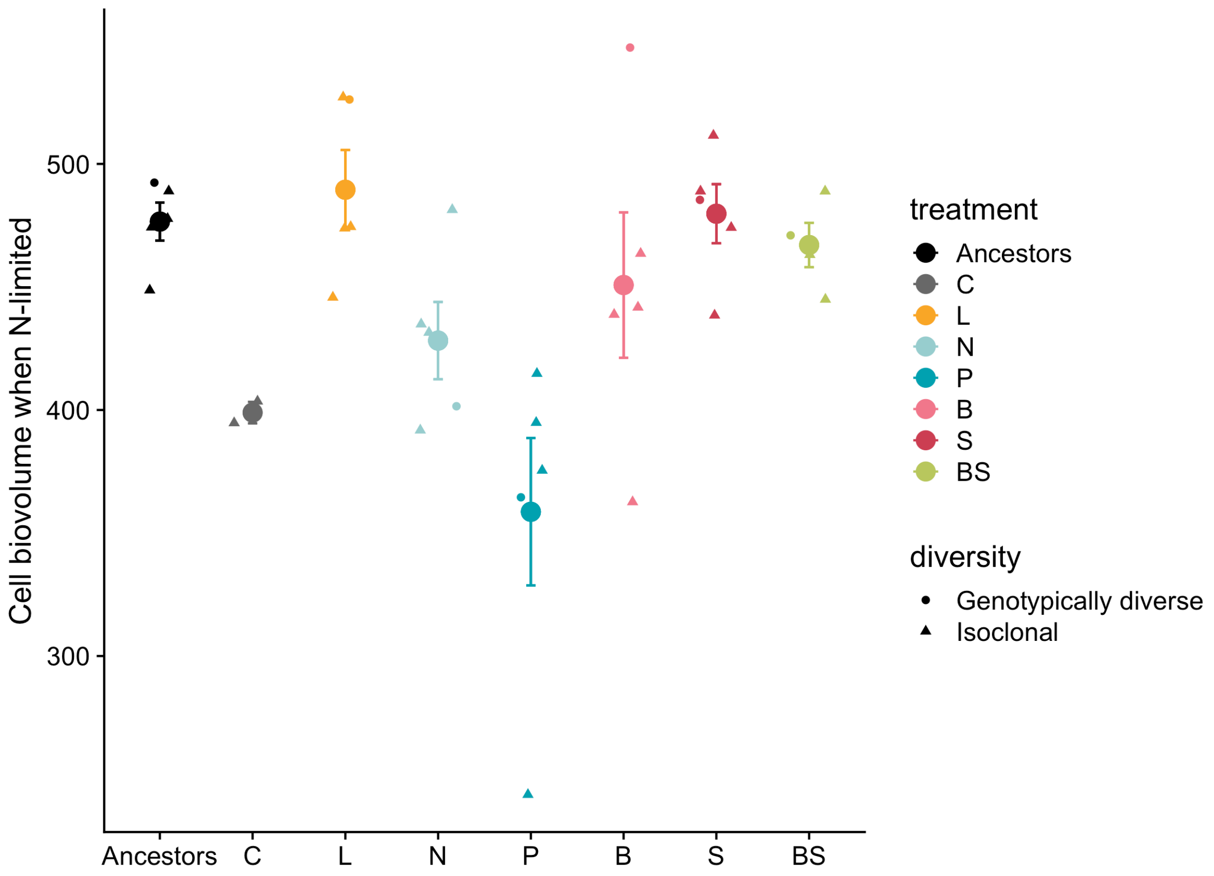


**Figure S8.** Cell biovolume ($um^3$) estimated in common garden conditions in batch culture in 10 uM N media. Colours correspond to the selection environment: A: ancestors, C: COMBO, L: light-limited, P: P-limited, N: N-limited, B: biotically depleted media, S: high salt, BS: biotically depleted and high salt.

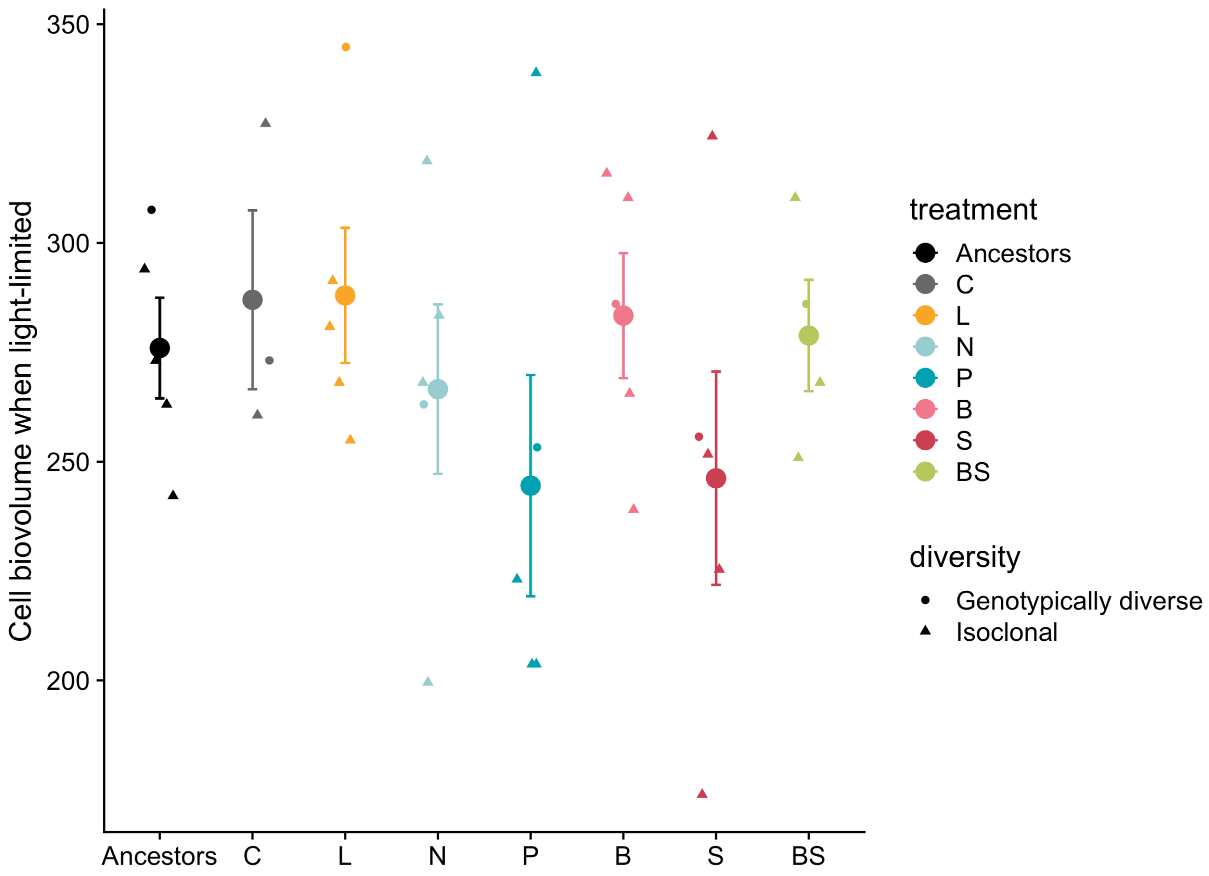


**Figure S9.** Cell biovolume (um$^3$) estimated in common garden conditions in batch culture in low light conditions (20 umol/m$^2$/s). Colours correspond to the selection environment: A: ancestors, C: COMBO, L: light-limited, P: P-limited, N: N-limited, B: biotically depleted media, S: high salt, BS: biotically depleted and high salt.

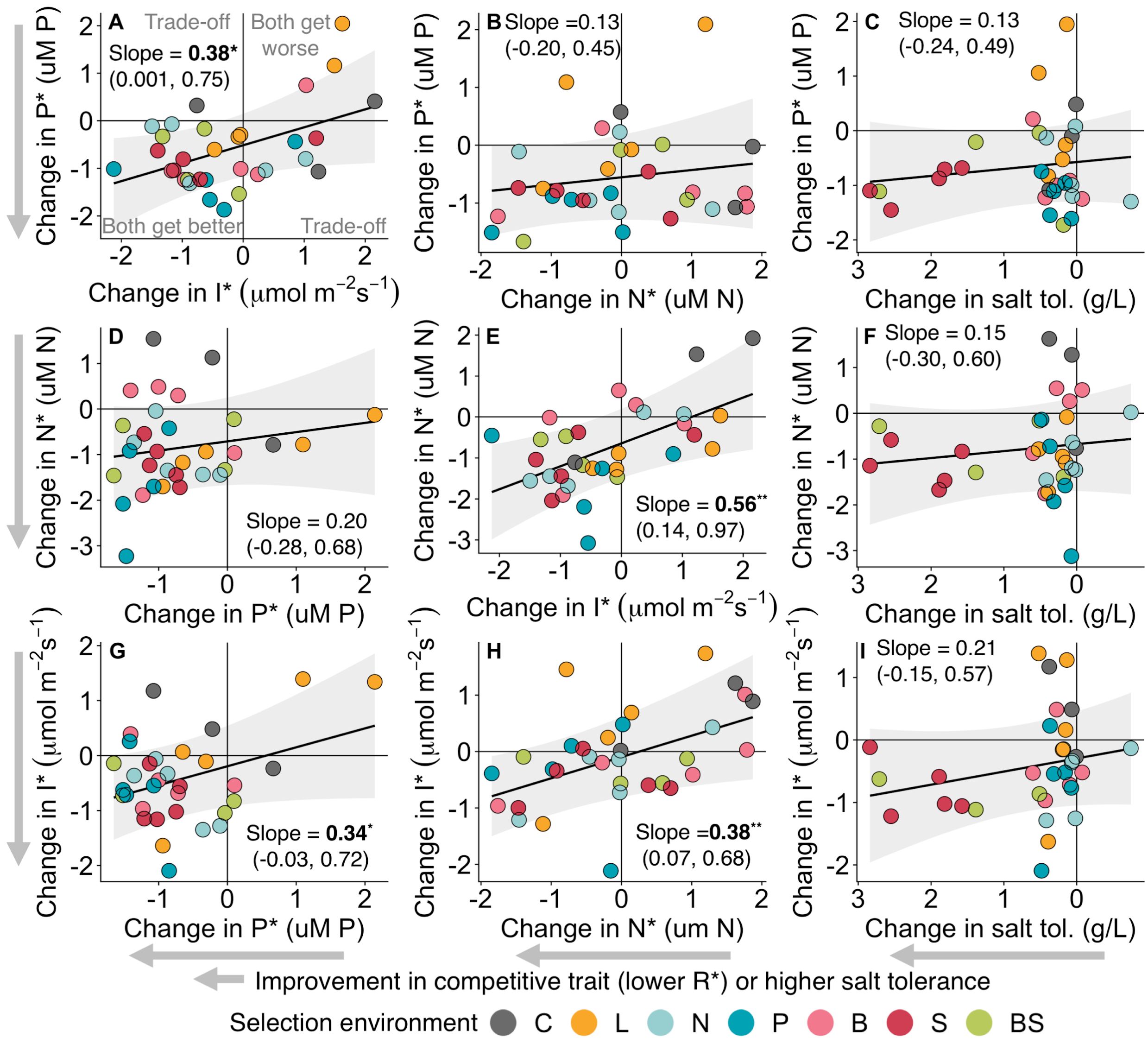


**Figure S10.** Partial regression plots showing how changes in two traits are related to each other, while holding all factors in the statistical model that are not being displayed constant (model results in ESM Tables 1-3). Negative slopes indicate a trade-off in trait changes, while positive slopes indicate positively associated trait changes. Note that the x-axis for salt tolerance is reversed in panels C, F and I to be consistent with the direction of adaptive trait change in the other panels. Grey arrows indicate the direction of improvement in competitive traits or salt tolerance. Panel A describes the four quadrants delineating areas where both traits get better, both get worse, or one gets better while the other gets worse (trade-off). Panels A, B and E are the same as in Figure 5 in the main text.

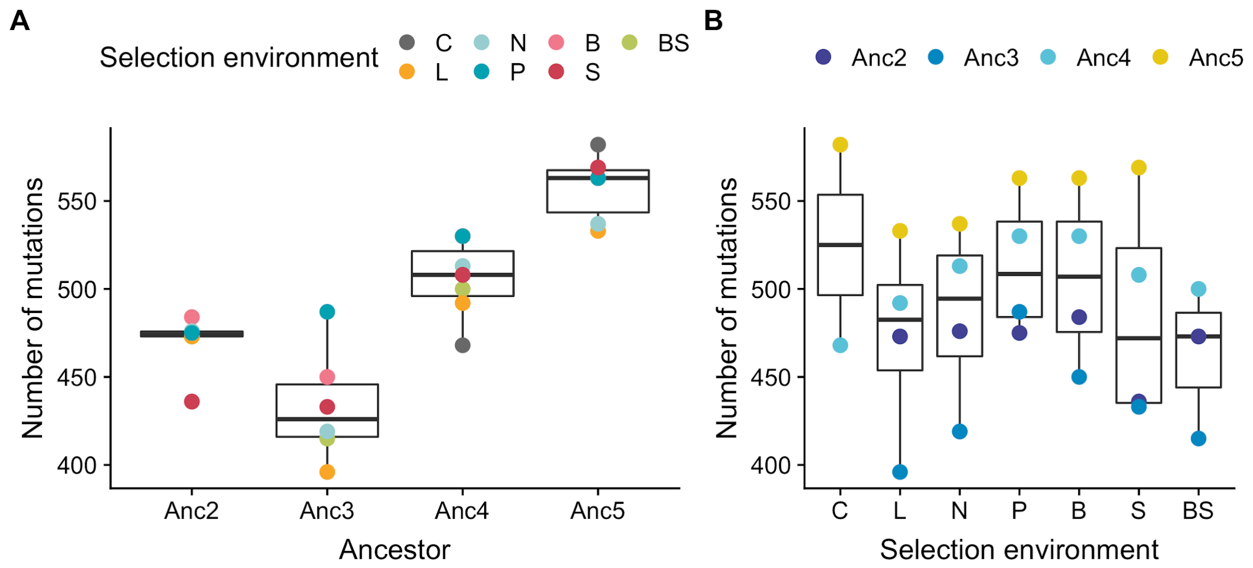


**Figure S11.** Number of mutations as a function of ancestry (A) or selection environment (B). The effect of ancestry on the number of mutations was highly significant (ANOVA p < 1e-7) while the selection environment had no significant effect (ANOVA p = 0.788).

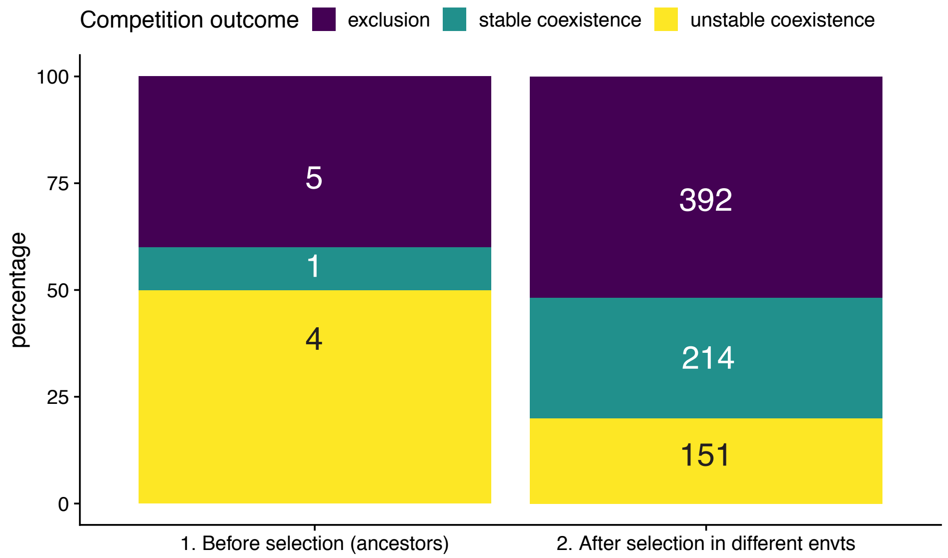


**Figure S12.** Percentage of all possible pairwise competitive interactions before (left side of panel) and after selection in seven different resource environments (right side of panel).

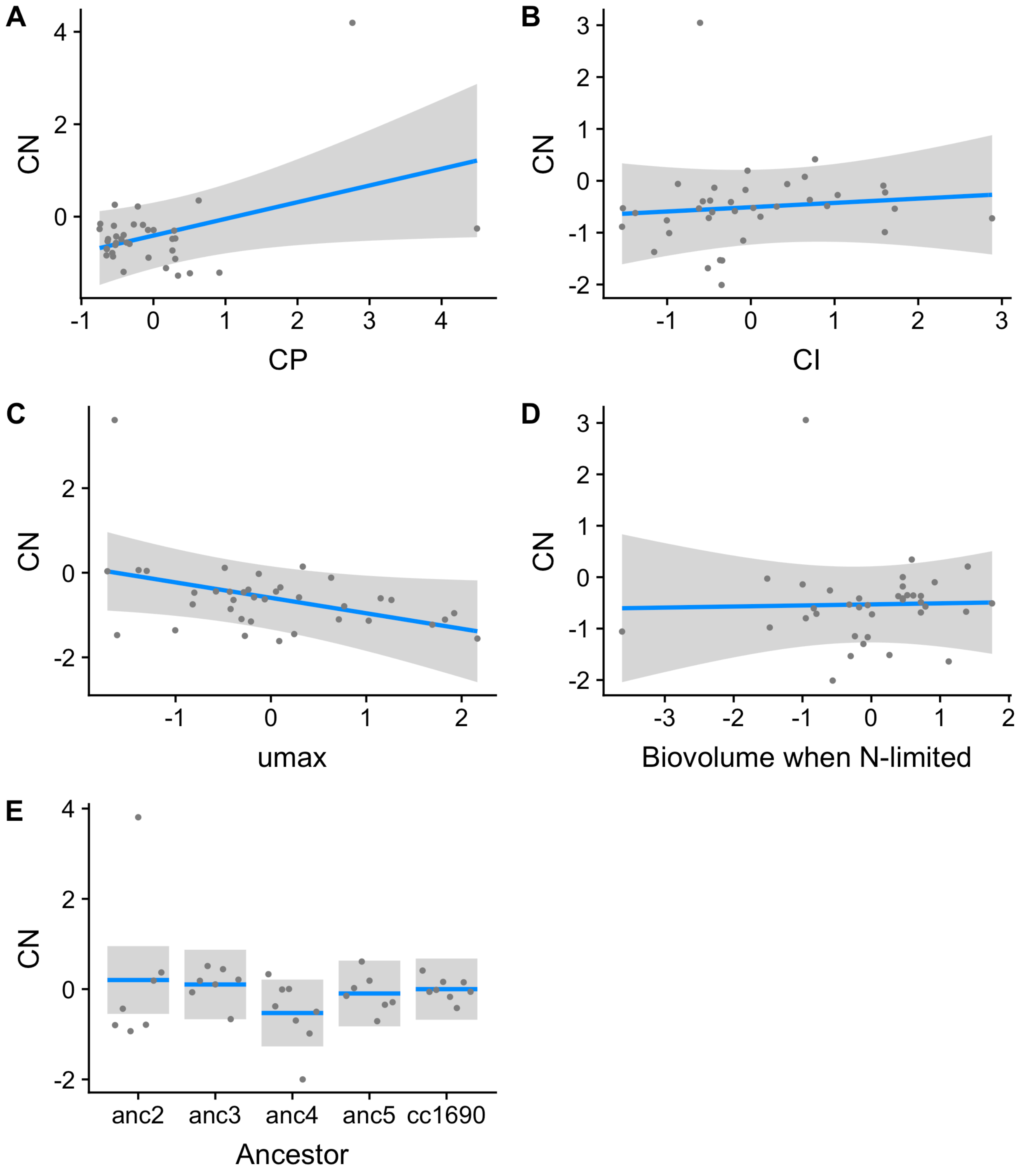


**Figure S13.** Partial regression plots showing how competitive ability for nitrogen (CN) changes as a function of A) competitive ability for phosphorus (CP), B) competitive ability for light (CI), C) umax (derived from fits of Monod curve over a gradient of nitrogen supply; Figure 4B), D) cell biovolume when growing in nitrogen limited conditions (ESM Figure S8) while holding all factors in the statistical model that are not being displayed constant (at their median value). For model results, see ESM Table 4.

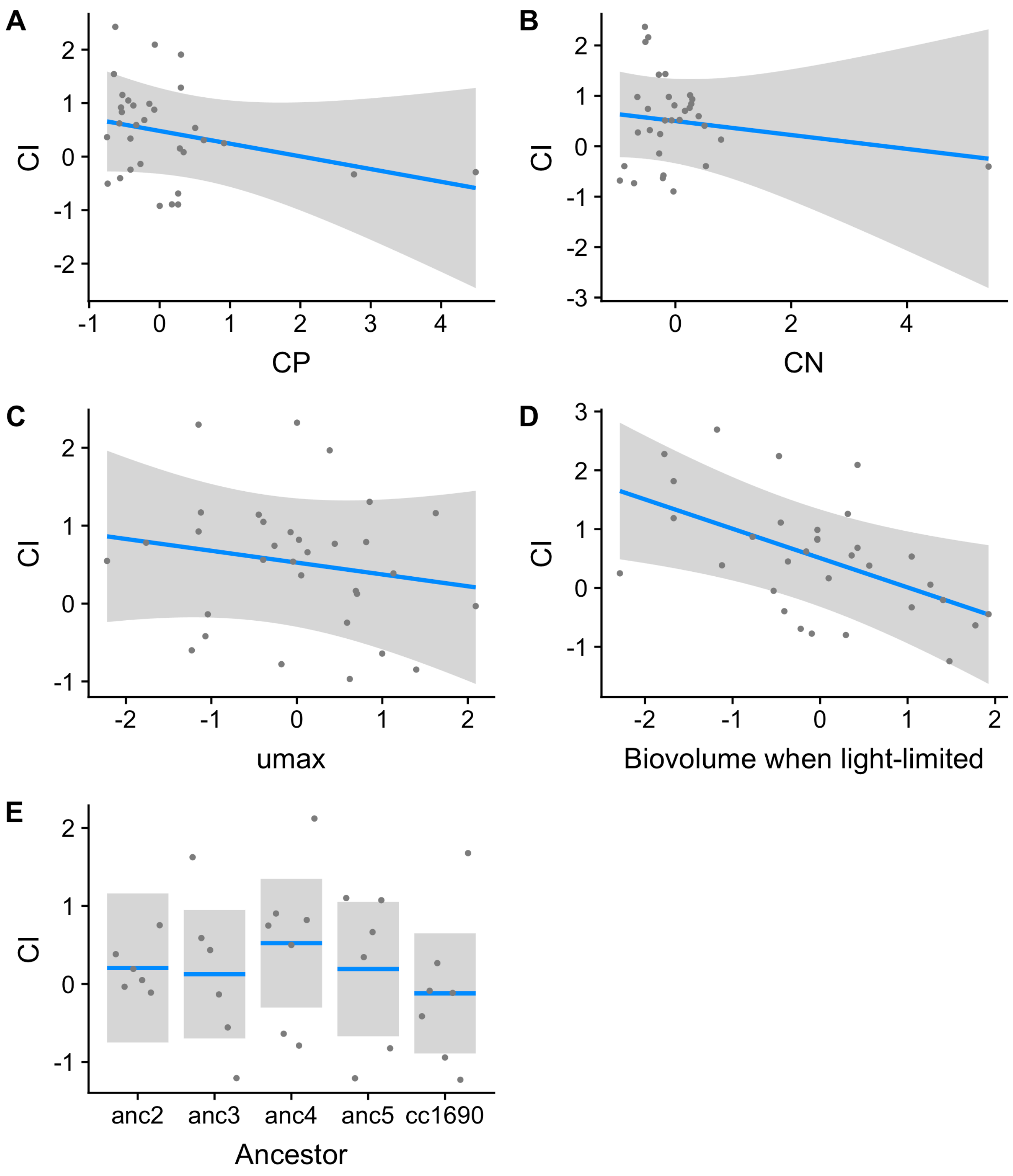


**Figure S14.** Partial regression plots showing how competitive ability for light (CI) changes as a function of A) competitive ability for phosphorus (CP), B) competitive ability for nitrogen (CN), C) umax (derived from fits of Monod curve over a gradient of light supply; Figure 4C), D) cell biovolume when growing in light limited conditions (ESM Figure S9) while holding all factors in the statistical model that are not being displayed constant (at their median value). For model results, see ESM Table 5.

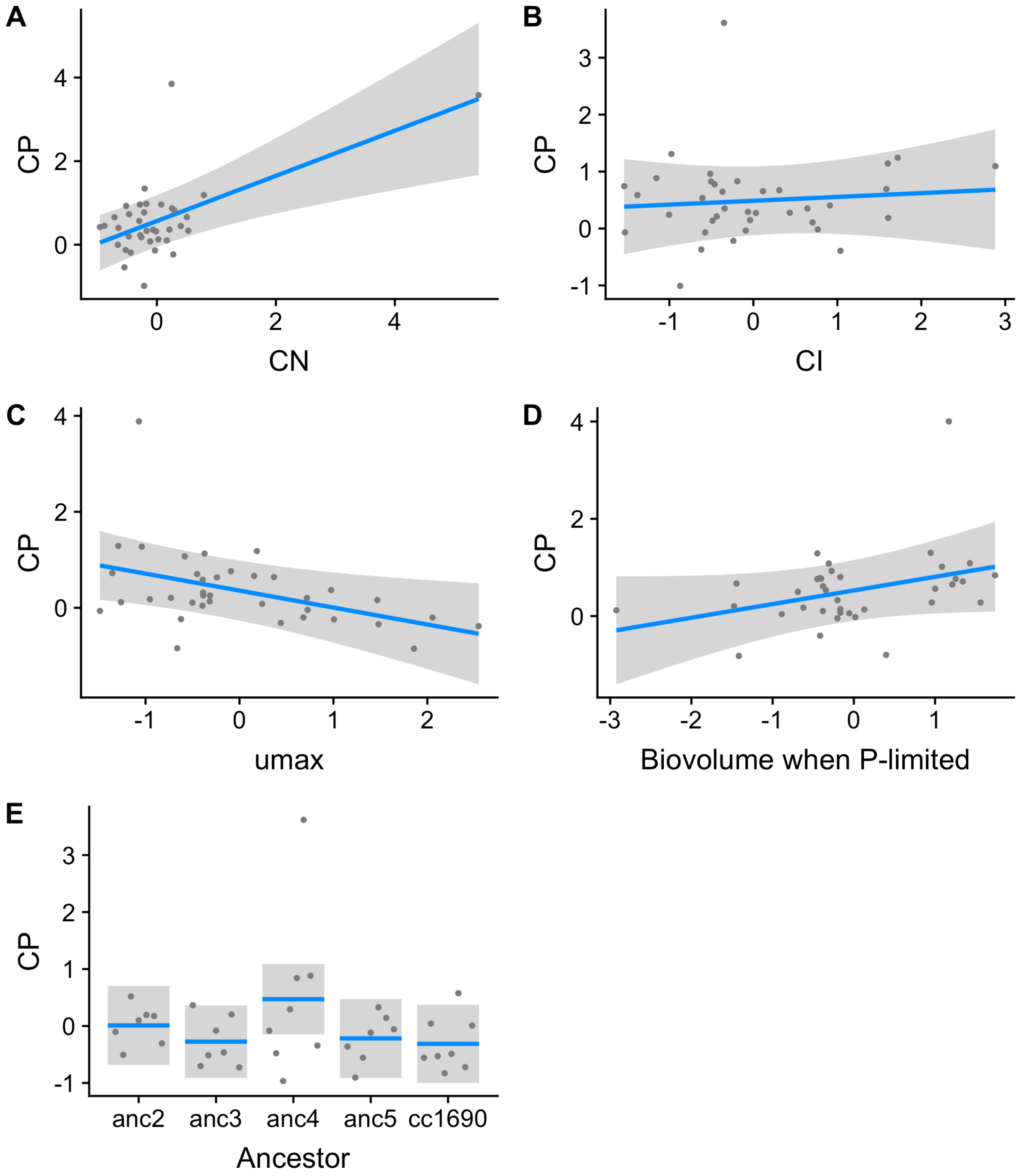


**Figure S15.** Partial regression plots showing how competitive ability for phosphorus (CP) changes as a function of A) competitive ability for nitrogen (CN), B) competitive ability for light (CI), C) umax (derived from fits of Monod curve over a gradient of phosphorus supply; Figure 4A), D) cell biovolume when growing in phosphorus limited conditions (ESM Figure S7) while holding all factors in the statistical model that are not being displayed constant (at their median value). For model results, see ESM Table 6.

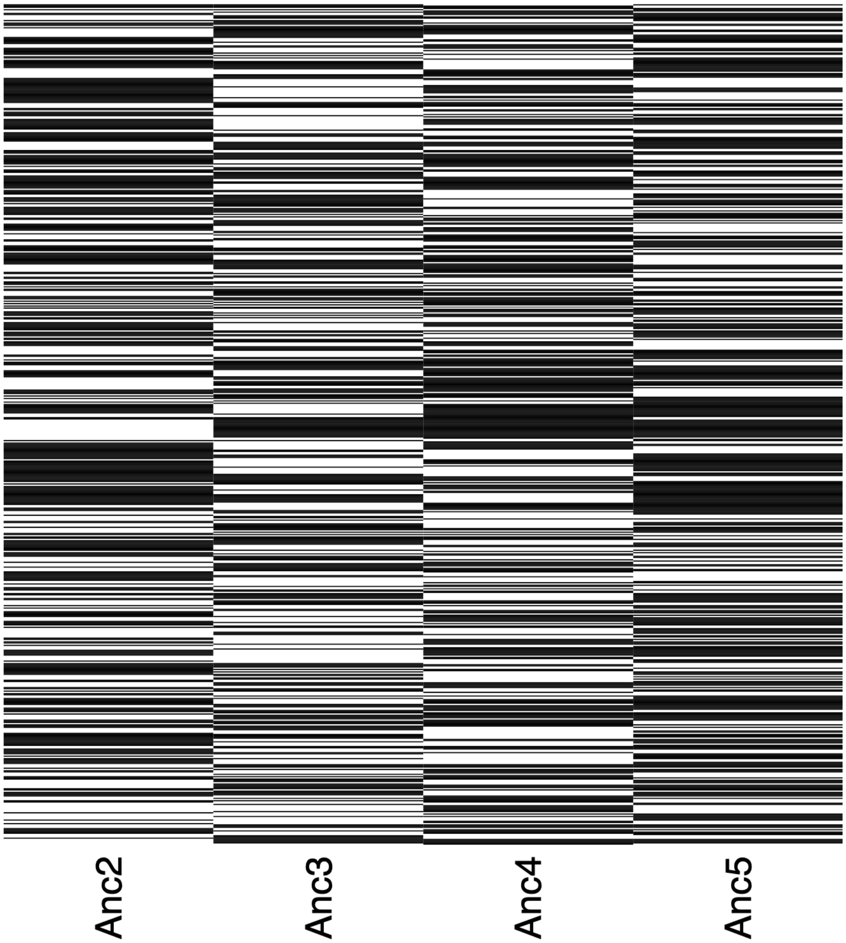


**Figure S16.** Variable loci between isolated from cc1690 population. Individual isolates are indicated on y-axis while the variable loci are on x-axis. Black indicates a SNP called against *C. reinhardtii* reference version 5.5. This figure demonstrates that the SNPs for each isolonal population against the published reference genome differ from one another.

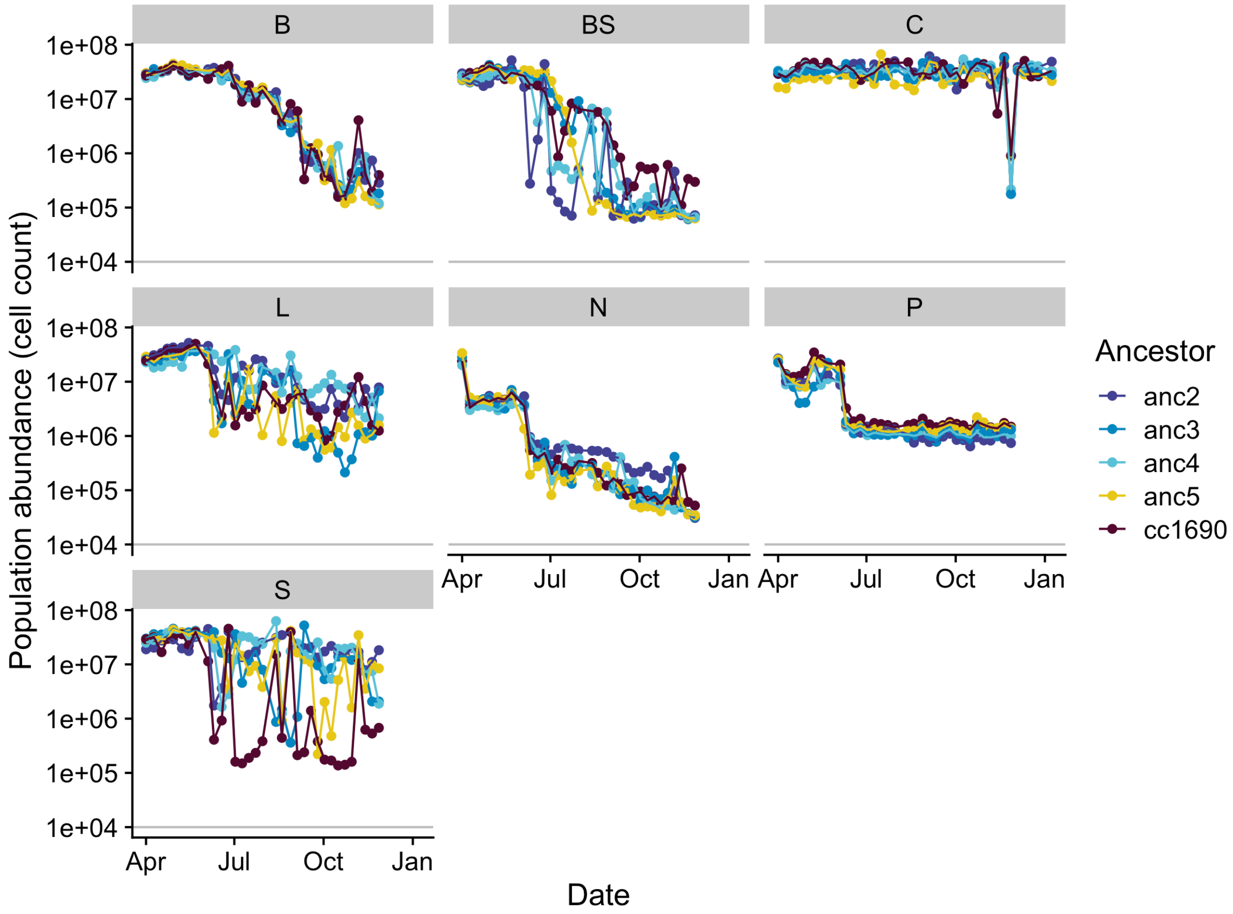


**Figure S17.** Population abundance trajectories over the evolution experiment estimated via Relative Fluorescence Units (RFU),converted to estimates of population abundance using flow cytometry. Note that the y-axis is on a log scale. Panels correspond to selection environments (B: biotically depleted, BS: Biotically-depleted + high salt, C: COMBO, L: light limitation, N: nitrogen limitation, P: phosphorus limitation, S: high salt).

### References:

1. Monod J. 1949 The growth of bacterial cultures. *Annu. Rev. Microbiol.* **3**, 371–394. (doi:10.1146/annurev.mi.03.100149.002103)

2. Grover JP. 1997 Resource Competition. (doi:10.1007/978-1-4615-6397-6)

3. Stearns SC. 1983 The Influence of Size and Phylogeny on Patterns of Covariation among Life-History Traits in the Mammals. *Oikos* **41**, 173–187.

(doi:10.2307/3544261)

4. Reich PB. 2014 The world-wide 'fast--slow'plant economics spectrum: a traits manifesto. *J. Ecol.* **102**, 275–301.

5. Eilers PHC, Peeters JCH. 1988 A model for the relationship between light intensity and the rate of photosynthesis in phytoplankton. *Ecol. Modell.* **42**, 199–215. (doi:10.1016/0304-3800(88)90057-9)
