## Appendix C :Supplementary Tables for "The evolution of competitive ability for essential resources"

### Electronic Supplementary Materials: Appendix C

#### Supplementary Tables

#### Tables 1-7

### The evolution of competitive ability for essential resources

Joey R. Bernhardt[1*], Pavel Kratina[2], Aaron Pereira[1], Manu Tamminen[3], Mridul K. Thomas[4], Anita Narwani[1]

[1]Aquatic Ecology Department, Eawag, Überlandstrasse 133, CH-8600 Dübendorf, Switzerland

[2]School of Biological and Chemical Sciences, Queen Mary University of London, Mile End Road, London E1 4NS, United Kingdom.

[3]Department of Biology, University of Turku, Natura, University Hill, 20014 Turku, Finland

[4]Centre for Ocean Life, DTU Aqua, Technical University of Denmark, Kongens Lyngby, Denmark

| | Change in I* | | |
|---|---|---|---|
| | (1) | (2) | (3) |
| **Change in salt tol** | -0.21 (-0.57, 0.15) | | |
| **Change in N*** | | $0.38^{**}$ (0.07, 0.68) | |
| **Change in P*** | | | $0.34^{*}$ (-0.03, 0.72) |
| **Change in size** | 0.11 (-0.25, 0.47) | 0.04 (-0.30, 0.37) | 0.10 (-0.24, 0.44) |
| **Anc 3** | -0.59 (-1.60, 0.41) | -0.66 (-1.58, 0.27) | -0.44 (-1.43, 0.55) |
| **Anc 4** | -0.35 (-1.30, 0.60) | 0.11 (-0.84, 1.07) | -0.29 (-1.20, 0.63) |
| **Anc 5** | $1.23^{**}$ (0.09, 2.36) | $1.35^{**}$ (0.31, 2.39) | $1.30^{**}$ (0.21, 2.39) |
| **cc1690** | -0.77 (-1.81, 0.27) | -0.64 (-1.59, 0.31) | -0.75 (-1.75, 0.25) |
| **Constant** | 0.07 (-0.69, 0.83) | -0.21 (-0.88, 0.47) | 0.11 (-0.61, 0.83) |
| Observations | 32 | 32 | 32 |
| $R^2$ | 0.46 | 0.54 | 0.49 |
| Adjusted $R^2$ | 0.33 | 0.43 | 0.37 |
| *Note:* | | | $^{*}$p<0.1; $^{**}$p < 0.05; $^{***}$p<0.01; (95% CI) |

**Table 1.** Multiple regression fits of change in I* of descendant populations relative to their ancestors, as a function of changes in cell biovolume (size; when growing in light limiting conditions (ESM Figure S9) and ancestry, as well as changes in salt tolerance (model 1), changes in N* (model 2), changes in P* (model 3).

| | Change in P* | | |
|---|---|---|---|
| | (1) | (2) | (3) |
| **Change in salt tol** | -0.13 (-0.49, 0.24) | | |
| **Change in I*** | | 0.38* (0.001, 0.75) | |
| **Change in N*** | | | 0.13 (-0.20, 0.45) |
| **Change in size** | 0.05 (-0.33, 0.43) | -0.08 (-0.45, 0.29) | 0.01 (-0.37, 0.39) |
| **Anc 3** | -0.58 (-1.60, 0.44) | -0.41 (-1.39, 0.57) | -0.63 (-1.64, 0.39) |
| **Anc 4** | -0.18 (-1.17, 0.81) | -0.12 (-1.05, 0.81) | -0.05 (-1.10, 1.01) |
| **Anc 5** | -0.23 (-1.45, 1.00) | -0.87 (-2.14, 0.41) | -0.23 (-1.44, 0.99) |
| **cc1690** | -0.05 (-1.01, 0.92) | 0.32 (-0.65, 1.28) | 0.04 (-0.94, 1.01) |
| **Constant** | -0.39 (-1.14, 0.36) | -0.40 (-1.09, 0.28) | -0.51 (-1.25, 0.23) |
| Observations | 32 | 32 | 32 |
| $R^2$ | 0.10 | 0.20 | 0.10 |
| Adjusted $R^2$ | -0.12 | 0.01 | -0.12 |
| *Note:* | | | *p<0.1; **p < 0.05; ***p<0.01; (95% CI) |

**Table 2.** Multiple regression fits of change in P* of descendant populations relative to their ancestors, as a function of changes in cell biovolume (size; when growing in phosphorus limited conditions (ESM Figure S7), ancestry and changes in salt tolerance (model 1), changes in I* (model 2), changes in N* (model 3).

| | Change in N* | | |
|---|---|---|---|
| | (1) | (2) | (3) |
| **Change in salt tol** | -0.15 (-0.60, 0.30) | | |
| **Change in I*** | | 0.56** (0.14, 0.97) | |
| **Change in P*** | | | 0.20 (-0.28, 0.68) |
| **Change in size** | 0.004 (-0.45, 0.46) | -0.09 (-0.48, 0.31) | -0.08 (-0.52, 0.37) |
| **Anc 3** | -0.06 (-1.28, 1.15) | 0.30 (-0.82, 1.42) | 0.03 (-1.21, 1.27) |
| **Anc 4** | -1.28** (-2.50, -0.06) | -1.15** (-2.23, -0.06) | -1.31** (-2.51, -0.10) |
| **Anc 5** | -0.68 (-1.90, 0.53) | -1.25** (-2.41, -0.08) | -0.61 (-1.83, 0.62) |
| **cc1690** | -0.53 (-1.70, 0.63) | -0.02 (-1.12, 1.09) | -0.50 (-1.66, 0.66) |
| **Constant** | 0.61 (-0.32, 1.54) | 0.48 (-0.31, 1.26) | 0.57 (-0.31, 1.46) |
| Observations | 32 | 32 | 32 |
| $R^2$ | 0.21 | 0.37 | 0.22 |
| Adjusted $R^2$ | 0.02 | 0.22 | 0.03 |
| *Note:* | | | *p<0.1; **p < 0.05; ***p<0.01; (95% CI) |

**Table 3.** Multiple regression fits of change in N* of descendant populations relative to their ancestors, as a function of changes in cell biovolume (size; when growing in nitrogen limited conditions (ESM Figure S8), ancestry, and changes in salt tolerance (model 1), changes in I* (model 2), changes in P* (model 3).

| | Competitive ability for nitrogen, CN (1/N*) |
|---|---|
| **CP** | 0.36** (0.02, 0.70) |
| **CI** | 0.08 (-0.26, 0.42) |
| **Biovolume** | 0.02 (-0.32, 0.36) |
| **umax** | -0.37* (-0.74, 0.01) |
| **Anc 3** | -0.10 (-1.12, 0.92) |
| **Anc 4** | -0.73 (-1.68, 0.22) |
| **Anc 5** | -0.30 (-1.29, 0.70) |
| **cc1690** | -0.20 (-1.17, 0.77) |
| **Constant** | 0.28 (-0.42, 0.97) |
| Observations | 37 |
| $R^2$ | 0.37 |
| Adjusted $R^2$ | 0.18 |
| *Note:* | *p<0.1; **p < 0.05; ***p<0.01; (95% CI) |

**Table 4.** Multiple regression fits of competitive ability for nitrogen (CN), as a function of competitive ability for phosphorus (CP), competitive ability for light (CI), cell biovolume when growing in nitrogen limited conditions (ESM Figure S8) and umax (derived from fits of Monod curve over a gradient of nitrogen supply; Figure 2B).

| | Competitive ability for light, CI (1/I*) |
|---|---|
| **CP** | -0.24 (-0.64, 0.16) |
| **CN** | -0.14 (-0.53, 0.26) |
| **Biovolume** | -0.50** (-0.87, -0.12) |
| **umax** | -0.15 (-0.52, 0.22) |
| **Anc 3** | -0.08 (-1.27, 1.11) |
| **Anc 4** | 0.32 (-0.83, 1.47) |
| **Anc 5** | -0.01 (-1.30, 1.27) |
| **cc1690** | -0.33 (-1.47, 0.82) |
| **Constant** | 0.12 (-0.74, 0.99) |
| **Observations** | 32 |
| **$R^2$** | 0.34 |
| **Adjusted $R^2$** | 0.11 |
| *Note:* | *p<0.1; **p < 0.05; ***p<0.01; (95% CI) |

**Table 5.** Multiple regression fits of competitive ability for light (CI), as a function of competitive ability for nitrogen (CN), competitive ability for light (CI), cell biovolume when growing in light limited conditions (ESM Figure S9) and umax (derived from fits of Monod curve over a gradient of light availability; Figure 2C).

| | Competitive ability for phosphorus, CP (1/P*) |
|---|---|
| CN | 0.54*** (0.24, 0.84) |
| CI | 0.07 (-0.25, 0.38) |
| Biovolume | 0.28 (-0.05, 0.61) |
| umax | -0.35** (-0.64, -0.07) |
| Anc 3 | -0.28 (-1.17, 0.60) |
| Anc 4 | 0.46 (-0.43, 1.35) |
| Anc 5 | -0.23 (-1.13, 0.67) |
| cc1690 | -0.32 (-1.21, 0.56) |
| Constant | 0.07 (-0.57, 0.70) |
| Observations | 37 |
| $R^2$ | 0.48 |
| Adjusted $R^2$ | 0.33 |
| *Note:* | *p<0.1; **p < 0.05; ***p<0.01; (95% CI) |

**Table 6.** Multiple regression fits of competitive ability for phosphorus (CP), as a function of competitive ability for nitrogen (CN), competitive ability for light (CI), cell biovolume when growing in phosphorus limited conditions (ESM Figure S7) and umax (derived from fits of Monod curve over a gradient of phosphorus supply; Figure 2A).

| | B | BS | C | L | N | P | S |
|---|---|---|---|---|---|---|---|
| Anc2 | 484 | 473 | NA | 473 | 476 | 475 | 436 |
| Anc3 | 450 | 415 | NA | 396 | 419 | 487 | 433 |
| Anc4 | 530 | 500 | 468 | 492 | 513 | 530 | 508 |
| Anc5 | 563 | NA | 582 | 533 | 537 | 563 | 569 |

**Table 7.** Number of variable SNPs between the ancestors and descendants from different selection environments: C: COMBO, L: light-limited, P: P-limited, N: N-limited, B: biotically depleted media, S: high salt, BS: biotically depleted and high salt.
